# RetroCHMP3 Blocks Budding of Enveloped Viruses Without Blocking Cytokinesis

**DOI:** 10.1101/2020.08.30.273656

**Authors:** Lara Rheinemann, Diane Miller Downhour, Kate Bredbenner, Gaelle Mercenne, Kristen A. Davenport, Phuong Tieu Schmitt, Christina R. Necessary, John McCullough, Anthony P. Schmitt, Sanford M. Simon, Wesley I. Sundquist, Nels C. Elde

## Abstract

Many enveloped viruses require the endosomal sorting complexes required for transport (ESCRT) pathway to exit infected cells. This highly conserved pathway mediates essential cellular membrane fission events, which restricts the acquisition of adaptive mutations to counteract viral co-option. Here, we describe duplicated and truncated copies of the ESCRT-III factor CHMP3 that block ESCRT-dependent virus budding and that arose independently in New World monkeys and mice. When expressed in human cells, these retroCHMP3 proteins potently inhibit release of retroviruses, paramyxoviruses, and filoviruses. Remarkably, retroCHMP3 proteins have evolved to reduce interactions with other ESCRT-III factors and to have little effect on cellular ESCRT processes, revealing routes for decoupling cellular ESCRT functions from viral exploitation. The repurposing of duplicated ESCRT-III proteins thus provides a mechanism to generate broad-spectrum viral budding inhibitors without blocking highly conserved essential cellular ESCRT functions.

## Introduction

The endosomal sorting complexes required for transport (ESCRT) pathway mediates essential cellular membrane fission events such as multivesicular body formation, cytokinetic abscission, and resealing of the post-mitotic nuclear envelope. Loss of ESCRT function is therefore highly detrimental in many cultured cell lines. Early-acting ESCRT factors are recruited to cellular sites of membrane fission by specific adapter proteins that, in turn, recruit core ESCRT-III charged multivesicular body proteins (CHMPs, *e.g.*, CHMP2, CHMP3, and CHMP4), which form membrane-associated heteropolymeric filaments. These ESCRT-III protein filaments then recruit and are remodeled by the AAA ATPase VPS4, which is required for membrane constriction and fission (Christ et al., 2017; Henne et al., 2013; McCullough et al., 2018; Scourfield and Martin-Serrano, 2017).

To facilitate budding from cellular membranes, the structural proteins of enveloped viruses, including the human immunodeficiency virus type 1 (HIV-1), contain ESCRT recruitment motifs, termed late assembly domains, that mimic the motifs employed by host ESCRT adapter proteins (Votteler and Sundquist, 2013). Such piracy can generate evolutionary pressure to select for mutations in host proteins that abrogate the interaction with viral proteins (Daugherty and Malik, 2012). In this case, however, mutations in ESCRT proteins that abrogate viral pathway recruitment would likely also interfere with essential cellular ESCRT functions and be selected against (Elde and Malik, 2009). Therefore, the host has limited potential for adaptations of the ESCRT pathway that prevent exploitation by viruses while still maintaining interactions necessary for cellular processes.

Retrocopies are gene duplications that arise by reverse transcription and genomic insertion of mRNAs by retrotransposons such as long interspersed nuclear elements (LINEs). In rare cases, retrocopies evolve new functions that differ from the parental gene (Kaessmann et al., 2009; Kubiak and Makalowska, 2017). Here, we describe the independent evolution of retrocopies of the ESCRT-III protein CHMP3 from New World monkey and mouse species that can potently block ESCRT-dependent virus budding. Remarkably, these retroCHMP3 proteins have evolved to spare cellular ESCRT functions in abscission, and therefore specifically counteract the hijacking of the ESCRT pathway by viruses.

## Results

### RetroCHMP3 proteins inhibit HIV-1 budding

We searched for CHMP3 retrocopies by manual BLAST-like Alignment Tool (BLAT) analyses of genomes in the University of California Santa Cruz (UCSC) genome browser (Kent, 2002) or by manual National Center for Biotechnology Information (NCBI) BLAST analyses (Altschul et al., 1990) of whole-genome shotgun contigs. Our searches identified two independent duplications of the gene encoding the ESCRT-III protein CHMP3 in ancestors of New World monkeys and house mice. These gene duplications appear to be LINE-1-mediated, as evidenced by the absence of introns and the presence of flanking target site duplications, which are hallmarks of LINE-1 retrotransposition (see **Table S1** for details). Importantly, the parental CHMP3 gene (∼222 amino acids) remains intact in all species, which presumably allowed the retrocopies to diverge in function. These retrocopies acquired independent premature stop codons, removing approximately 70 carboxy-terminal amino acids from the predicted protein product, in two New World monkeys; Guyanese squirrel monkey (*Saimiri sciureus sciureus*, retroCHMP3^SS^, 155 amino acids) and black-handed spider monkey (*Ateles geoffroyi*, retroCHMP3^AG^, 157 amino acids), and in house mice (*Mus musculus*, retroCHMP3^MM^, 147 amino acids, **Fig. 1A**). We verified the expected retroCHMP3 sequences by genomic DNA sequencing (**Table S1**, **Fig. S1A**, see Methods for details). RetroCHMP3 genes predicated to express similar truncated proteins have also been detected in other species, and are described elsewhere (Rheinemann et al., 2021).

**Fig. 1:**
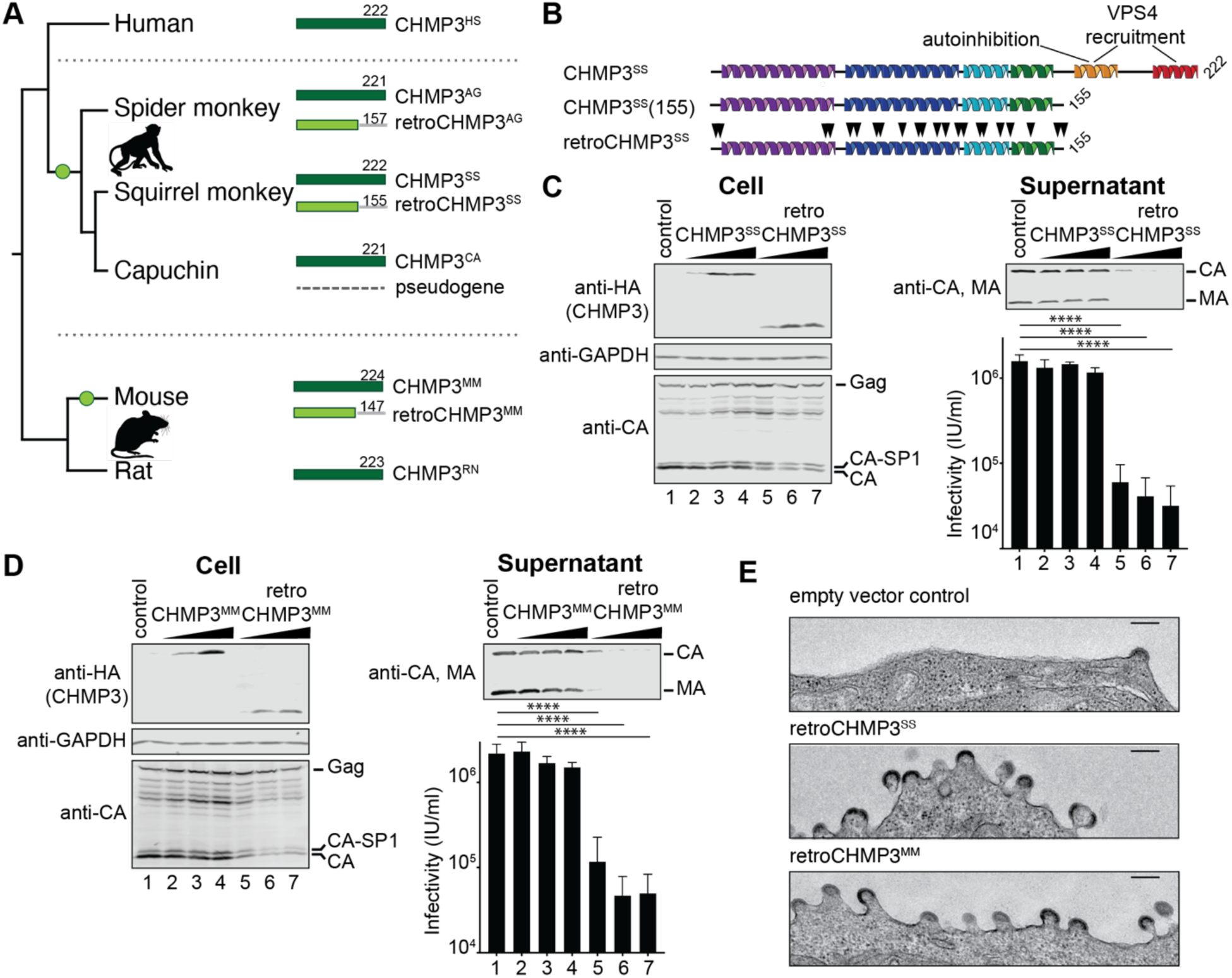
RetroCHMP3 proteins inhibit HIV-1 budding. **(A)** Schematic representation of the C-terminally truncated retrocopies of CHMP3 (light green) found in squirrel monkey (retroCHMP3^SS^), spider monkey (retroCHMP3^AG^), and mouse (retroCHMP3^MM^). Green circles indicate the acquisition of retrocopies. Parental full-length genes (dark green) are also present in all species. The CHMP3 retrocopy in capuchin is homologous to retroCHMP3^SS^, but degraded. **(B)** Schematic representation of full-length, parental squirrel monkey CHMP3 (CHMP3^SS^), C-terminally truncated parental CHMP3 (CHMP3^SS^(155)), and retroCHMP3^SS^. Black arrowheads indicate amino acid substitutions. **(C), (D)** Transient overexpression of HA-tagged retroCHMP3^SS^ **(C)** and retroCHMP3^MM^ **(D)** in human embryonic kidney (HEK293T) cells inhibits the release of HIV-1 proteins (CA, MA) and reduces viral titers in the supernatant. Titer graphs show mean ± SD from 3 experimental replicates. **** p<0.0001, *** p<0.001 by one-way ANOVA followed by Tukey’s multiple comparisons test. For a complete set of pairwise comparisons, see Table S3. **(E)** Transmission electron micrographs of cells transfected with an HIV-1 expression vector and retroCHMP3^SM^ or retroCHMP3^MM^. Scale bar 100nm.

A truncated human CHMP3 construct lacking the C-terminal 72 amino acids can potently inhibit HIV-1 budding (Zamborlini et al., 2006). This truncated construct presumably functions as a dominant-negative inhibitor of ESCRT pathway functions because it retains the core filament assembly domain but lacks the C-terminal elements required for autoinhibition and recruitment of VPS4 and its filament depolymerization activity (Bajorek et al., 2009; Lata et al., 2008; Muziol et al., 2006; Shim et al., 2007; Stuchell-Brereton et al., 2007) (**Fig. 1B**). We hypothesized that the naturally occurring retroCHMP3 proteins might therefore also exhibit antiviral activity, and we tested this idea by co-expressing either HA-tagged retroCHMP3^SS^ (**Fig. 1C**), retroCHMP3^MM^ (**Fig. 1D**), or retroCHMP3^AG^ (**Fig. S1B**) together with a proviral HIV-1 expression construct in human embryonic kidney (HEK293T) cells. In all three cases, retroCHMP3 expression potently inhibited virus release, as measured by reductions in the levels of viral MA (matrix) and CA (capsid) proteins released into the culture supernatant and by reductions in infectious titers. RetroCHMP3 expression also caused two additional phenotypes characteristic of HIV-1 budding defects or delays: 1) accumulation of the Gag processing intermediate, CA-SP1, relative to fully processed CA protein (Gottlinger et al., 1991), and 2) concomitant accumulation of budding virions seen at the plasma membrane by thin-section transmission electron microscopy (**Fig. 1E**). Therefore, retroCHMP3 proteins encoded by three different species can potently inhibit HIV-1 budding, presumably by dominantly inhibiting ESCRT pathway functions.

### RetroCHMP3 proteins broadly inhibit ESCRT-dependent budding

To test whether retroCHMP3 inhibition can be generalized to other ESCRT-dependent viruses, we co-expressed either retroCHMP3^SS^ or retroCHMP3^MM^ with proviral expression constructs for Friend murine leukemia virus (FrMLV, **Fig. 2A, S2A**) or equine infectious anemia virus (EIAV, **Fig. 2B, S2B**) in HEK293T cells. Both of these retroviruses have previously been shown to require the ESCRT pathway for budding (Garrus et al., 2001; Strack et al., 2003). RetroCHMP3 proteins reduced the release of FrMLV MA and CA proteins and viral titers in the supernatant without significantly altering the expression of viral proteins in the cell (**Fig. 2A, S2A**). EIAV titers were also reduced by retroCHMP3 protein expression (**Fig. 2B, S2B**), but in this case, the amount of EIAV CA protein in the supernatant was not substantially reduced. These observations are similar to our previous report that inhibiting ESCRT functions by depleting the ESCRT-III protein CHMP4B during EIAV budding also reduced viral titers but did not decrease virion-associated CA protein release (Sandrin and Sundquist, 2013). In that report, we also observed that EIAV virions released in the absence of CHMP4B exhibited aberrant tubular or multi-lobed virion morphologies, suggesting that Gag polymerization or bud neck closure defects led to the release of aberrant, non-infectious virions. In the case of retroCHMP3^SS^ co-expression, we similarly observed a large number of immature EIAV virions that were either budding or were closely associated with the cell surface, and a subset displayed aberrant morphologies (**Fig. 2C**, black arrowheads). Hence, retroCHMP3 inhibition resembled the phenotype seen for depletion of an essential ESCRT-III protein during EIAV budding, although the number of tubular extracellular EIAV virions was larger in the absence of CHMP4B than in the presence of retroCHMP3.

**Fig. 2:**
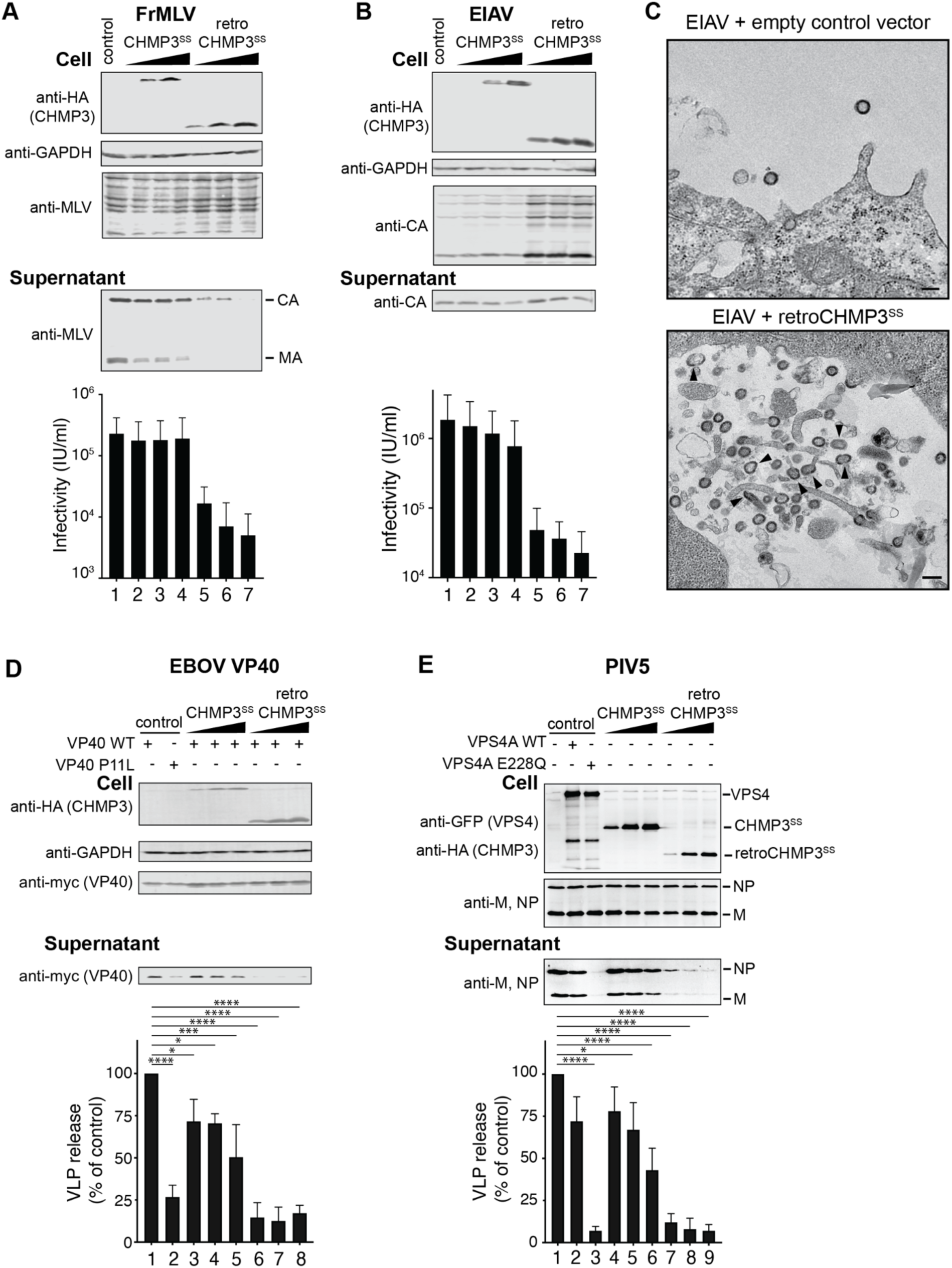
RetroCHMP3 proteins broadly inhibit ESCRT-dependent budding. **(A), (B)** Transient overexpression of HA-tagged retroCHMP3^SS^ in HEK293T cells reduces the release of viral proteins and viral titers in the supernatant of Friend murine leukemia virus (FrMLV, **A**) and equine infectious anemia virus (EIAV, **B**). Titer graphs show mean ± SD from 3 experimental replicates. **(C)** Transmission electron micrographs of cells transfected with EIAV expression vectors and empty control vector or retroCHMP3^SS^. Black arrowheads point to tubular and multi-lobed virions. Scale bar 100nm. **(D), (E)** Transient overexpression of HA-tagged retroCHMP3^SS^ in HEK293T cells reduces the release of Ebola virus VP40 protein **(D)** and parainfluenza virus 5 VLPs **(E)** into the supernatant. Western blots show representative results from 3 experimental replicates. Bar graphs show integrated intensities of viral protein bands in supernatant normalized to cellular fraction from 3 experimental replicates. **** p<0.0001, *** p<0.001, * p<0.05 by one-way ANOVA followed by Tukey’s multiple comparisons test. Only comparisons with the empty vector control (lane 1) are shown for clarity. For a complete set of pairwise comparisons, see Table S3.

We also tested the inhibitory effect of retroCHMP3 proteins on the budding of an ESCRT-dependent filovirus (Ebola virus, EBOV, **Fig. 2D, S2C**) and an ESCRT-dependent paramyxovirus (parainfluenza virus 5, PIV 5, **Fig. 2E, S2D**) (Martin-Serrano et al., 2001; Schmitt et al., 2005). In both cases, retroCHMP3^SS^ and retroCHMP3^MM^ proteins inhibited virion release. Specifically, release of virus-like particles (VLPs) formed by the EBOV VP40 protein was inhibited by both retroCHMP3 proteins but not by parental full-length CHMP3 proteins. The effect was comparable to that of a positive control mutation in the VP40 late assembly domain (VP40 P11L) that abrogates ESCRT recruitment (Martin-Serrano et al., 2004). Release of PIV5 VLPs produced by the co-expression of the PIV5 M, NP, and HN proteins was similarly reduced by the retroCHMP3 proteins. This effect was comparable to the positive control for loss of ESCRT activity, in this case induced by overexpression of a dominant-negative mutant of VPS4 (VPS4A E228Q). Taken together, these data demonstrate that retroCHMP3 proteins can broadly block the release of highly divergent virus families by inhibiting ESCRT functions during budding.

### RetroCHMP3 proteins evolved to lose cytotoxicity

The ESCRT pathway performs a series of important cellular functions, including cytokinetic abscission, and loss of ESCRT function is therefore typically cytotoxic. Indeed, overexpression of parental CHMP3 proteins that were artificially truncated to the same length as retroCHMP3 proteins dramatically reduced cell viability after 24 hours, consistent with dominant-negative inhibition of ESCRT functions by these constructs (**Fig. 3A, S3A, S3B**). Strikingly, however, overexpression of retroCHMP3^SS^ only very mildly reduced cell viability and was nearly as non-toxic as overexpression of the full-length, parental CHMP3^SS^ protein (**Fig. 3A**). RetroCHMP3^SS^ acquired 22 amino acid substitutions in comparison to the truncated parental CHMP3^SS^(155) (black arrowheads, **Fig. 1B**), suggesting that these amino acid substitutions, or a subset of them, cause the reduced cytotoxicity of retroCHMP3^SS^. RetroCHMP3^MM^ and retroCHMP3^AG^ also displayed lower cytotoxicity than their C-terminally truncated parental CHMP3 counterparts. In these cases, however, the proteins were moderately more cytotoxic than the corresponding full-length CHMP3 proteins (**Figs. S3A, B**). Accordingly, cellular expression of HIV-1 Gag, which was also assessed after 24 hours and is dependent on cell viability, was reduced moderately in samples transfected with retroCHMP3^MM^ (**Fig. 1C**) and retroCHMP3^AG^ (**Fig. S1B**), but not retroCHMP3^SS^ (**Fig. 1B**). RetroCHMP3^MM^ and retroCHMP3^AG^ acquired 12 and 14 amino acid substitutions, respectively, compared to their truncated parental proteins. Notably, none of the retroCHMP3 proteins share common mutations (**Fig. S1A)**, demonstrating that different combinations of amino acid substitutions can lead to a reduction of cytotoxicity.

**Fig. 3:**
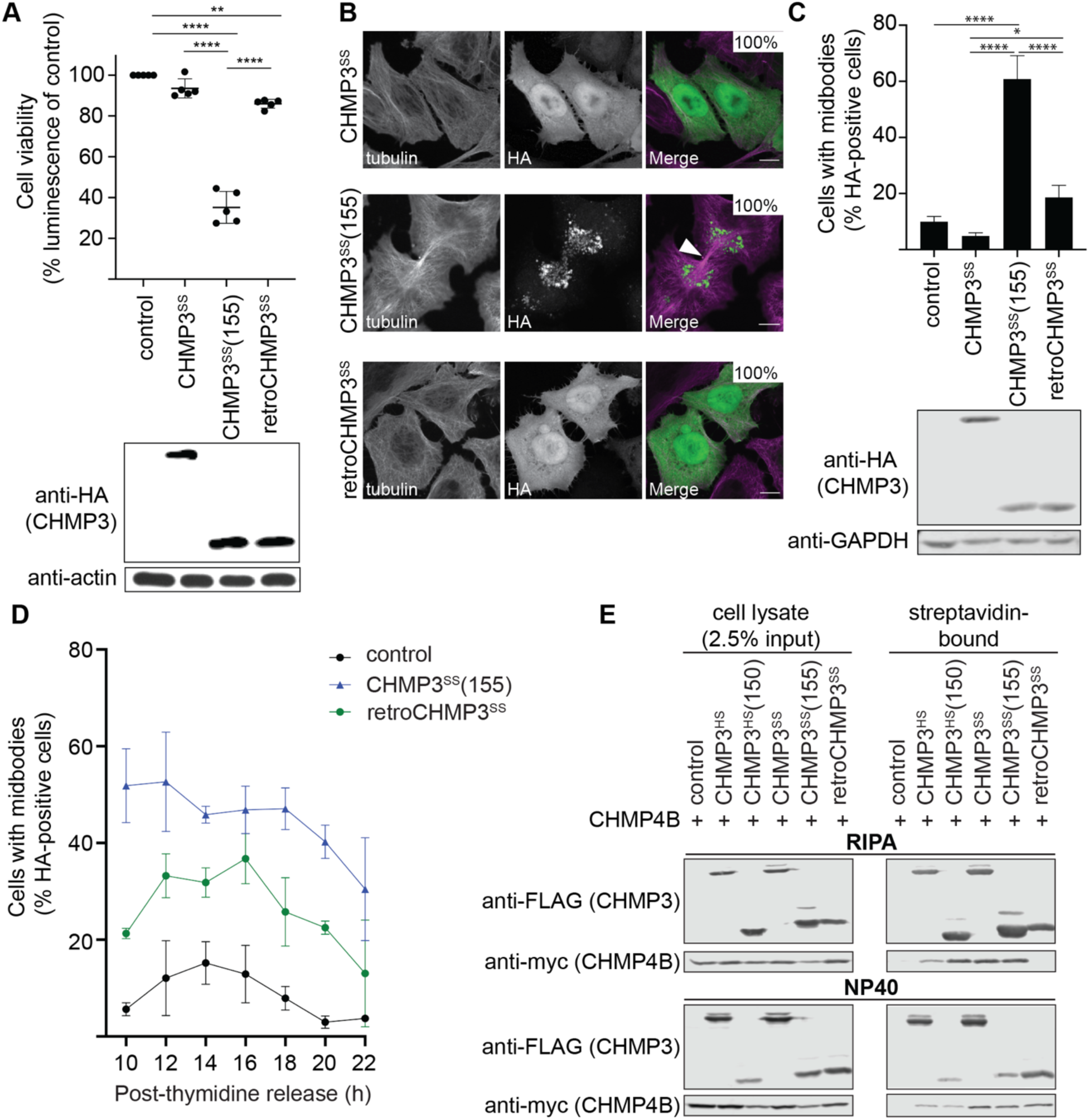
RetroCHMP3^SS^ evolved to lose cytotoxicity. **(A)** Viability of HEK293T cells transiently transfected with HA-tagged CHMP3^SS^ variants at 24 hours after transfection, as determined in a luminescent ATP cell viability assay. Mean ± SD from 5 experimental replicates with n≥ 6 each. **(B)** Immunofluorescence showing the dispersed (CHMP3^SS^ and retroCHMP3^SS^) and punctate (CHMP3^SS^(155)) subcellular localization phenotypes in HeLa N cells. Scale bar 10 μm. Insets show percentages of cells with the two depicted phenotypes (diffuse or punctate: n≥ 450 cells/condition). White arrowhead points to a midbody. **(C)** Number of midbody-linked cells in unsynchronized cultures transiently transfected with the designated HA-tagged CHMP3^SS^ variants 20 h after transfection. Mean ± SD from 3 experimental replicates, each with 150 HA-positive cells/condition. **(D)** Number of midbody-linked cells in synchronized cultures transiently transfected with the designated HA-tagged CHMP3^SS^ variants. Times are post-thymidine release. Mean ± SD from 3 experimental replicates with > 290 HA-positive cells total/timepoint. **(E)** Pulldown of myc-tagged human CHMP4B with STrEP/FLAG-tagged human CHMP3 (CHMP3^HS^), C-terminally truncated human CHMP3 (CHMP3^HS^(150)), or CHMP3^SS^ variants. Cell lysis and pulldowns were performed in RIPA or NP-40 buffer, as indicated. Representative Western blot from 3 experimental repeats. **** p<0.0001, ** p<0.01, * p<0.05 by one-way ANOVA followed by Tukey’s multiple comparisons test comparisons with p>0.05 omitted for clarity.

Cell imaging experiments revealed that cytotoxicity levels correlated with the subcellular localization patterns of the different CHMP3 proteins (**Fig. 3B**): Specifically, the two non-toxic constructs, parental CHMP3^SS^ and retroCHMP3^SS^, dispersed homogeneously throughout the cytoplasm, with some enrichment in the nucleus. In contrast, the toxic CHMP3^SS^(155) protein formed perinuclear punctae. Similarly, non-toxic parental CHMP3^MM^ was dispersed, whereas CHMP3^MM^(147) showed a punctate localization pattern (**Fig. S3C**). RetroCHMP3^MM^ displayed a mixed pattern with dispersed and punctate staining, consistent with incomplete detoxification. These data suggest that C-terminally truncated parental CHMP3 proteins formed insoluble aggregates due to the lack of the autoinhibitory and VPS4 recruitment regions, but that additional amino acid substitutions in retroCHMP3^SS^ and retroCHMP3^MM^ allowed these proteins to fully or partially restore their solubility to that of their respective parental CHMP3 proteins.

To test whether the cytotoxicity of the different CHMP3 proteins could be explained by ESCRT-dependent cell division phenotypes, we examined the effects of the different constructs on midbody resolution during cytokinetic abscission. Both truncated and retroCHMP3 proteins were recruited to the midbody, as was the full-length parental protein **(Fig. S3D)**, in good accord with a previous report (Dukes et al., 2008). Consistent with their relative cellular toxicities, and in line with previously observed negative effects of C-terminally truncated human CHMP3 on cytokinesis (Dukes et al., 2008), we found that CHMP3^SS^(155) overexpression dramatically increased the number of unresolved midbodies as compared to either CHMP3^SS^ or retroCHMP3^SS^ **(Fig. 3C).** Midbody numbers were much lower in the presence of retroCHMP3^SS^, but were still slightly elevated over the control construct, consistent with the slight cytotoxicity observed for this construct (**Fig. 3A**). To determine the effect of retroCHMP3^SS^ on the kinetics of abscission, we synchronized cells by addition of thymidine, which arrests cells in the G1/S phase. Release of the thymidine block then allows for synchronous progression through the remaining phases of the cell cycle. Cultures transfected with an empty control plasmid accumulated cells connected by midbodies until a peak was reached around 14 hours after thymidine release, after which the number of unresolved midbodies dropped because most cells had completed abscission at 20 hours after thymidine release (**Fig. 3D**). Cells transfected with CHMP3^SS^(155) had elevated numbers of unresolved midbodies at all timepoints, consistent with inhibition of midbody resolution already seen in asynchronous cultures. Cultures transfected with retroCHMP3^SS^ also exhibited elevated midbody numbers, and the peak of accumulation was delayed by about two hours relative to control cells. Eventually, the number of midbodies dropped as cells completed abscission. These observations suggest that midbodies are eventually resolved in cells transfected with retroCHMP3^SS^, albeit more slowly than in control cells. Thus, both the defects in abscission and reductions in cell viability were much more severe for CHMP3^SS^(155) than for retroCHMP3^SS^ and CHMP3^SS^, suggesting that abscission defects are a major driver of cytotoxicity.

We also tested how retroCHMP3 affected another ESCRT-dependent cellular process; degradation of the epidermal growth factor receptor (EGFR). In response to stimulation with EGF, EGFR is ubiquitylated at the plasma membrane, sorted into intraluminal vesicles (ILVs) at endosomes in an ESCRT-dependent manner, and ultimately delivered to lysosomes for degradation (Eden et al., 2009; Raiborg and Stenmark, 2009). Overexpression of retroCHMP3^SS^ delayed the EGF-mediated degradation of EGFR to the same extent as overexpression of a well-characterized inhibitor of the ESCRT pathway, an ATPase-deficient mutant of VPS4A, VPS4A E228Q (**Fig. S3E**). Therefore, in contrast to abscission, which was inhibited to a lesser extent by retroCHMP3^SS^ compared to the dominant-negative CHMP3^SS^(155), sorting of plasma membrane proteins into ILVs was equally affected by retroCHMP3^SS^ and the dominant-negative VPS4A E228Q construct. These observations suggest that multiple ESCRT processes are impaired by retroCHMP3, albeit with differential susceptibilities.

We next sought to determine how amino acid mutations functionally differentiate retroCHMP3^SS^ from CHMP3^SS^(155). The punctae formed by CHMP3^SS^(155) suggested that this engineered construct might inhibit the ESCRT pathway by co-polymerizing with other ESCRT-III partners and thereby functioning as a dominant-negative inhibitor of the pathway. We therefore examined the interaction of retroCHMP3^SS^ with the ESCRT-III protein CHMP4B, a major CHMP3 binding partner (Effantin et al., 2013). In pulldown experiments, full-length human CHMP3 (CHMP3^HS^) and full-length CHMP3^SS^ bound CHMP4B under both stringent (RIPA buffer, **Fig. 3E**, top panel) and mild (NP-40 buffer, **Fig. 3E**, bottom panel) conditions. C-terminally truncated CHMP3^HS^(150) and CHMP3^SS^(155) also bound strongly to CHMP4B under both conditions. In contrast, retroCHMP3^SS^ pulled down CHMP4B in NP-40 buffer, but not in RIPA buffer. This result demonstrates that mutations present in retroCHMP3^SS^ substantially weaken interactions with CHMP4B, perhaps by inhibiting formation of stable hetero-copolymeric filaments that remain intact under the more stringent RIPA buffer binding conditions (Bajorek et al., 2009; McCullough et al., 2015). Although the reduction in CHMP4B binding can explain why retroCHMP3^SS^ toxicity is attenuated, it leaves open the question of how retroCHMP3^SS^ can still inhibit ESCRT functions during virus budding.

### RetroCHMP3 alters CHMP4B and VPS4A recruitment to HIV-1 Gag assembly sites

To examine the effects of retroCHMP3^SS^ on the behavior of the ESCRT machinery at sites of virion assembly and budding, we used total internal reflection fluorescence microscopy (TIRF) to visualize how mEGFP-labeled Gag-Pol proteins co-assembled with mCherry-labeled versions of the essential late-acting ESCRT factors CHMP4B and VPS4A. (**Fig. 4**). This subviral construct includes the native Gag-Pol ribosomal frameshift site and therefore produces Gag-mEGFP and Gag-Pol-mEGFP proteins at normal viral ratios. The mEGFP fluorophore is inserted between the MA and CA domains of Gag so that both Gag and Gag-Pol proteins are fluorescently labeled, and the GFP is flanked by duplicated MA/CA cleavage sites for the viral protease (PR). Thus, assembling VLPs have the structural Gag protein and the enzymatic activities associated with Gag-Pol (including PR), and both Gag and Gag-Pol can undergo normal proteolytic maturation.

**Fig. 4:**
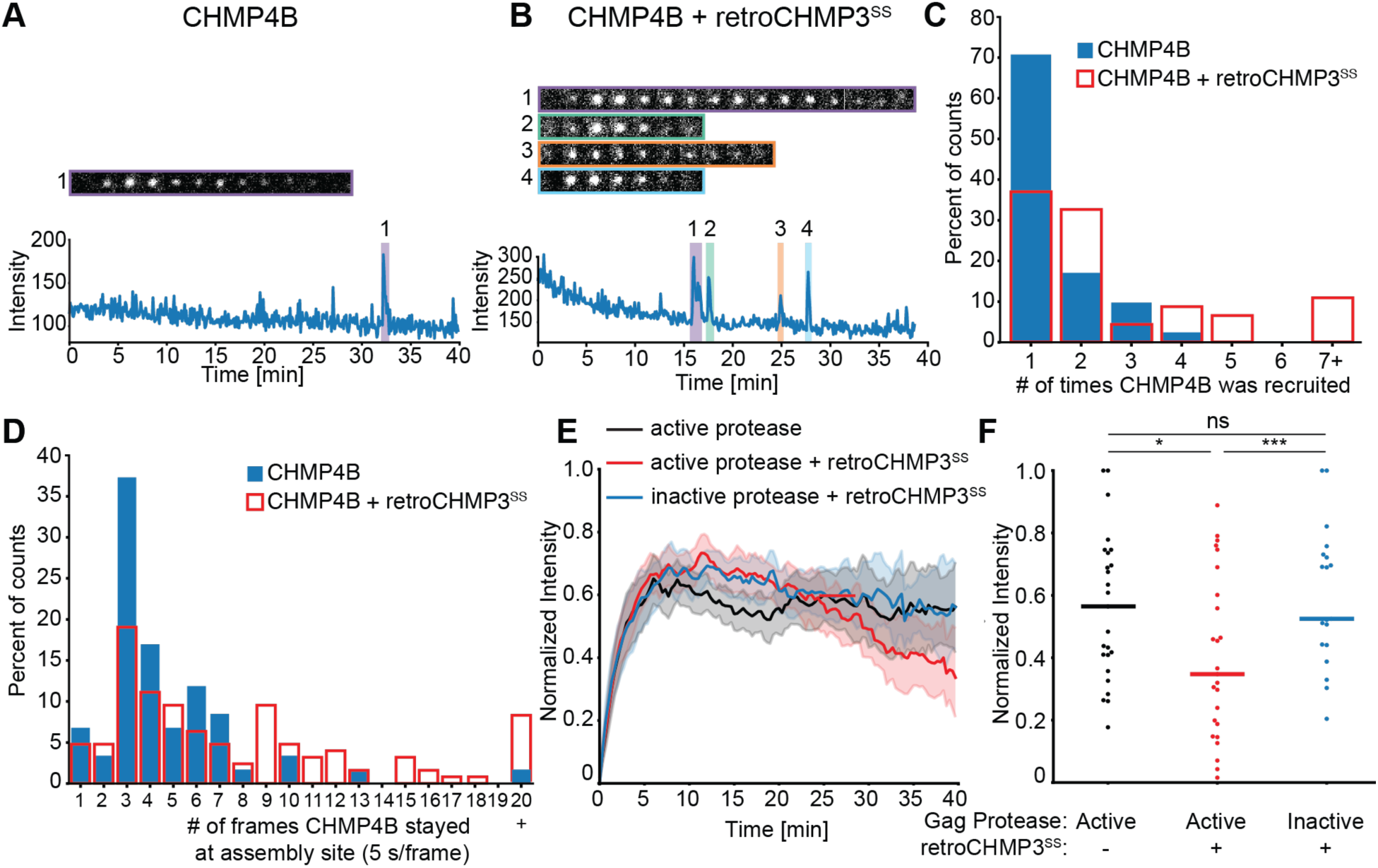
RetroCHMP3 alters CHMP4B recruitment to HIV-1 Gag assembly sites. **(A), (B)** Sample trace of mCherry-CHMP4B recruitment to Gag-Pol-mEGFP assembly sites in the absence **(A)** or presence **(B)** of retroCHMP3^SS^. Each numbered shaded region highlights one recruitment event and the corresponding sequence of images from that recruitment. The width of each image is 1.27 μm. **(C)** Histogram showing the number of times that CHMP4B was recruited to assembly sites in the absence (n= 46 assemblies) or presence (n=41 assemblies) of retroCHMP3^SS^. p<0.001 by one-tailed T-test. **(D)** Histogram showing the duration of each CHMP4B recruitment in the absence (n = 126 recruitments from 46 assemblies) or presence (n = 59 recruitments from 41 assemblies) of retroCHMP3^SS^. p<0.005 by one-tailed T-test. **(E)** Overlaid plots of average Gag recruitment for Gag-Pol-mEGFP without (black) or with (red) retroCHMP3^SS^ and Gag-Pol-mEGFP-D25A with retroCHMP3 ^SS^ (blue). The D25A mutation inactivates protease function. For each plot, solid line indicates average and shaded overlay represents the 95% confidence interval. **(F)** Plot of average final intensity of Gag-Pol-mEGFP for the final three frames for each trace that visibly recruited mCherry-CHMP4B in the absence (black) or presence (red) of retroCHMP3^SS^ or Gag-Pol-mEGFP-D25A with retroCHMP3^SS^ (blue). Colored bars indicate median. n = 25 assemblies per condition. *** p<0.001, * p<0.05 by two-tailed T-test. ns – not significant.

We and others previously reported that in a majority of VLP assembly events, ESCRT-III proteins and VPS4 are recruited once immediately prior to productive VLP budding, although multiple rounds of recruitment were observed in a minority of cases (Baumgartel et al., 2011; Bleck et al., 2014; Johnson et al., 2018; Jouvenet et al., 2011). Likewise, we observed that in the absence of retroCHMP3^SS^, mCherry-CHMP4B was recruited only once to mEGFP-marked virion assembly sites in 70% of the cases (**Fig. 4A, C**). In the presence of retroCHMP3^SS^, however, CHMP4B was recruited and lost multiple times at more than 60% of individual virion assembly sites (**Fig. 4B, C**). In extreme cases, more than seven cycles of CHMP4B recruitment to the same virion assembly site were observed. RetroCHMP3^SS^ also increased the duration of each cycle of recruitment of CHMP4B to the VLP (**Fig. 4D**). In the absence of retroCHMP3^SS^, 83% of CHMP4B recruitment events lasted between 5 and 30 seconds, with occasional longer intervals. In contrast, when retroCHMP3^SS^ was present, half of the visualized recruitment events lasted at least 30 seconds. These behaviors are consistent with multiple unproductive waves of ESCRT recruitment and failed membrane fission (Johnson et al., 2018). Thus, retroCHMP3^SS^ interferes with productive ESCRT-III filament assembly and function.

In complementary experiments, we found that retroCHMP3^SS^ did not alter the frequency of mCherry-VPS4A recruitment (**Fig. S4A-C**) but did reduce the residence time of VPS4A at the VLP, with nearly half of all VPS4A recruitments lasting ≤5 seconds in the presence of retroCHMP3^SS^ (**Fig. S4D**). Thus, retroCHMP3^SS^ also reduced the ability of ESCRT-III assemblies to bind and retain VPS4A, again suggesting that ESCRT-III filament assembly is compromised in the presence of retroCHMP3^SS^.

RetroCHMP3^SS^ also altered the behavior of Gag-Pol-mEGFP virion assembly (**Fig. 4E**). Under control conditions, Gag-Pol-mEGFP fluorescence rose sharply during the initial virion assembly phase and then remained at a stable plateau until the endpoint of the experiment. In contrast, when retroCHMP3^SS^ was present, Gag-Pol-mEGFP fluorescence at each VLP steadily decreased after reaching the initial plateau, so that the average mEGFP fluorescence intensity during the last three frames of each trace was lowered significantly (**Fig. 4F**). During viral assembly and maturation, the internal mEGFP fluorophore is released from Gag and Gag-Pol by PR processing. We found that the decrease in mEGFP-fluorescence required a functional protease because a construct with an inactive protease (Gag-Pol-D25A) did not lose fluorescence over time, even when retroCHMP3^SS^ was present (**Fig. 4E, F**). Previous work has shown that precise temporal control of the sequential steps of virion assembly and budding is essential for viral infectivity (Bendjennat and Saffarian, 2016). In particular, when budding is delayed by ESCRT pathway inactivation, PR auto-activation and Gag-Pol processing precede budding, the processed Gag and Gag-Pol components escape through the open bud neck, and viral infectivity is reduced. A similar process appears to be occurring in the presence of retroCHMP3^SS^, where protease activation during delayed budding liberates mEGFP to diffuse back into the cytoplasm. Taken together, these results suggest that retroCHMP3^SS^ impairs ESCRT-III filament assembly and delays membrane fission during budding.

### Expression of mouse retroCHMP3 can be induced by interferon signaling

To determine whether retroCHMP3 is expressed in mice and squirrel monkeys, we first surveyed a panel of squirrel monkey and mouse cell lines as well as mouse tissues (**Fig. S5**). We observed detectable levels of retroCHMP3 RNA in several mouse and squirrel monkey cell lines as well as mouse testes and heart, although RNA abundance appeared to be low under basal conditions. Intriguingly, the murine retroCHMP3 ORF contains candidate upstream promoter elements, including two putative interferon-responsive STAT1/STAT3 binding sites, and a murine retroviral related sequence (MuRRS) element (**Fig. 5A**). Endogenous retroviruses are known to encode elements that can regulate expression through interferon-induced signaling (Chuong et al., 2017), suggesting that retroCHMP3^MM^ might be regulated through immune signaling.

**Fig. 5:**
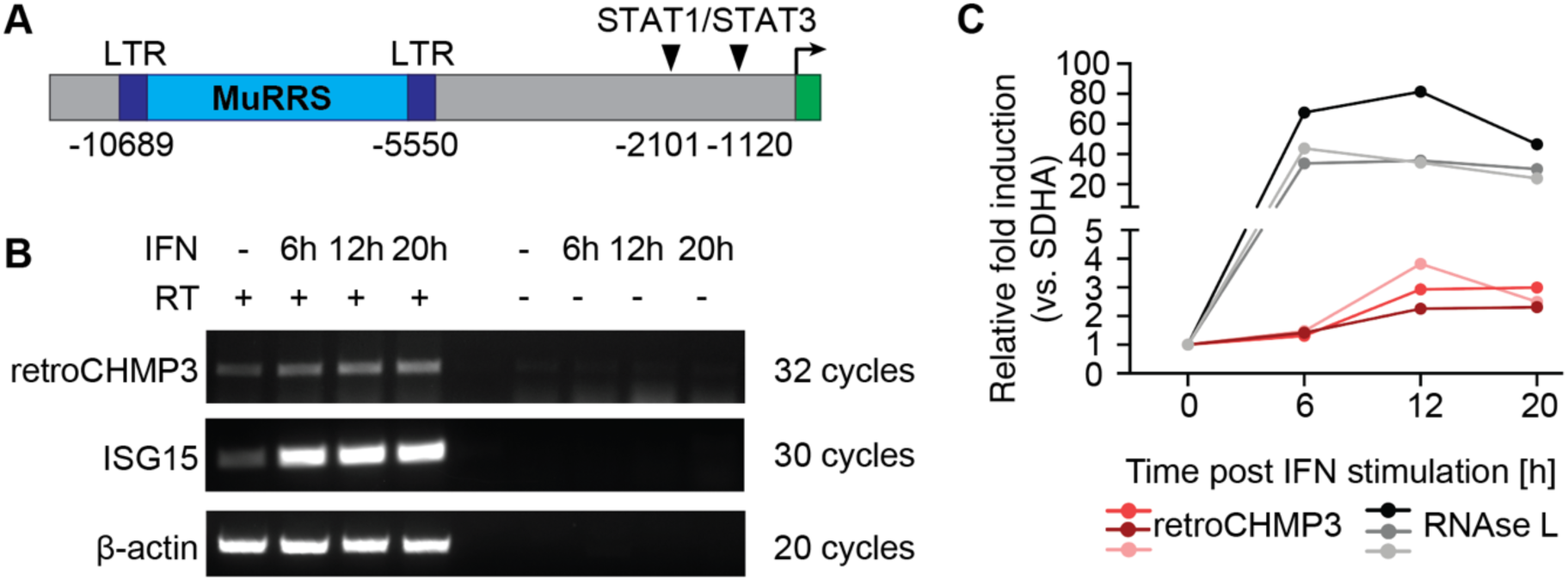
Expression of mouse retroCHMP3 may be controlled by interferon signaling. **(A)** Schematic of putative regulatory elements upstream of retroCHMP3 ORF in Mus musculus identified by transcription factor binding site prediction tools. Numbers indicate nucleotides relative to retroCHMP3 start codon. LTR – long terminal repeat, MuRRS – murine retrovirus-related sequence, ERV – endogenous retrovirus. **(B)** RT-PCR detection of retroCHMP3 RNA in mouse cardiac endothelial cells (MCECs) with and without interferon stimulation. RetroCHMP3 bands were excised, cloned, and sequenced to verify a match with the retroCHMP3 sequence. Representative gel from three independent biological repeats. RT – reverse transcriptase. **(C)** Droplet digital PCR (ddPCR) detection of retroCHMP3 (red) and RNAse L (grey, positive control) RNA in mouse cardiac endothelial cells (MCECs) with and without interferon stimulation. The housekeeping gene succinate dehydrogenase complex, subunit A (SDHA) was used for normalization. Each line represents one independent biological repeat.

Based on our observation of retroCHMP3^MM^ RNA expression in the mouse heart, we treated mouse cardiac endothelial cells with interferon and determined RNA expression levels by qRT-PCR (**Fig. 5B**) and droplet digital PCR (**Fig. 5C**). Interferon treatment modestly induced retroCHMP3^MM^ expression over 20 hours, compared to low levels of expression under basal conditions. RetroCHMP3 RNA levels in squirrel monkey B cells were not increased by interferon treatment (**Fig. S5B**). These results support a model in which the retrotransposed retroCHMP3^MM^ coding sequence landed in a region of the genome where it could be regulated by immune signaling.

## Discussion

We have identified retroCHMP3 genes in three different mammalian species, each of which independently gained a premature stop codon that removed ∼70 C-terminal amino acids. The frequency and retention of these independently evolved genes strongly suggest that they confer a selective advantage. When overexpressed in human cells, retroCHMP3 proteins potently block budding of a broad range of ESCRT-dependent viruses, yet they do not drastically inhibit the ESCRT-dependent cellular pathway of cytokinetic abscission. Their broad antiviral activity is consistent with the idea that retroCHMP3 proteins act by impeding a cellular function that is hijacked by a variety of viruses rather than targeting a specific viral protein.

A model explaining how retroCHMP3 can inhibit virus budding, yet differ from genetically engineered truncations of CHMP3 is shown in Fig. 6. Previous work has shown that truncated parental CHMP3 proteins can potently inhibit HIV-1 budding (Zamborlini et al., 2006), and also inhibit cellular ESCRT functions (Dukes et al., 2008). In this simple case, truncation activates the CHMP3 proteins for stable heteropolymeric filament formation, retains tight binding to CHMP4B (and possibly also other ESCRT-III proteins), and promotes cytoplasmic punctae formation. Truncated CHMP3 proteins could thus sequester ESCRT-III subunits away from functional sites such as the midbody and, even when recruited to the correct sites of action, create inert filaments that cannot undergo the VPS4-dependent subunit turnover required for membrane fission (Mierzwa et al., 2017). In contrast, retroCHMP3 proteins have evolved to reduce CHMP4B binding and apparently no longer form highly stable, inert ESCRT-III heteropolymers. We hypothesize that retroCHMP3 proteins therefore impair functional ESCRT-III filament assembly to a lesser degree than truncated CHMP3, thereby slowing down ESCRT-mediated membrane fission rather than blocking it entirely.

**Fig. 6:**
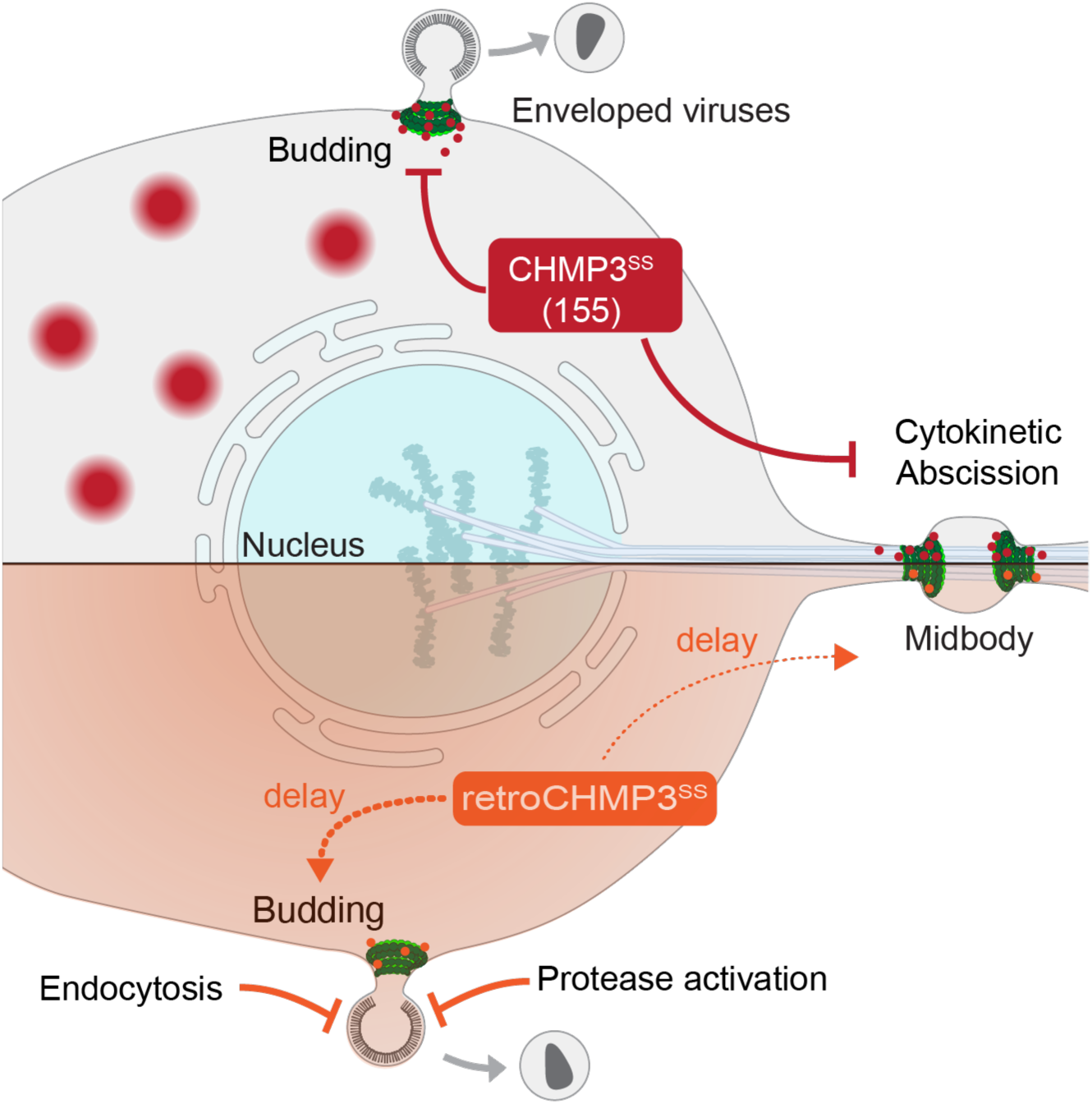
Models for truncated CHMP3 protein inhibition of ESCRT-mediated fission events and for retroCHMP3 detoxification.

We envision that retroCHMP3 proteins achieve selective antiviral activity without undue cellular toxicity by exerting a disproportionate impact on the rapid and time-sensitive process of virus budding. After the initial assembly of virions at the plasma membrane, ESCRT-III proteins and VPS4 are recruited transiently for seconds or minutes, followed by membrane fission (Baumgartel et al., 2011; Bleck et al., 2014; Johnson et al., 2018). HIV-1 assembly, budding, and maturation are highly coordinated processes, and budding delays that disturb the proper sequence thus also reduce infectivity by allowing processed viral components to escape from the virion prior to bud neck closure (Bendjennat and Saffarian, 2016). Furthermore, virion particles that remain attached to the plasma membrane can be reinternalized and degraded by endocytosis (Neil et al., 2006). In contrast, at least some essential cellular ESCRT-mediated processes, such as abscission, occur over much longer time scales of 1-2 hours (Elia et al., 2011; Guizetti et al., 2011; Hesse et al., 2012; Mackay and Ullman, 2015) and can tolerate very significant delays (Nahse et al., 2017), thereby sparing them from the most detrimental effects of retroCHMP3. In line with this, EGF-mediated degradation of the EGFR receptor, which displays similar dynamics as virus budding (Wenzel et al., 2018), is susceptible to inhibition by retroCHMP3 proteins. However, inhibition of receptor degradation presumably contributes less to acute cytotoxicity than does inhibition of cytokinesis, and therefore can be better tolerated.

Inhibition of ESCRT activity during infection has been described in other contexts: Intracellular bacteria such as *Mycobacteria* and *Salmonella* species replicate in membrane-bound vacuoles after their uptake into cells by phagocytosis or receptor-mediated endocytosis but eventually gain access to the host cytosol by breaching endolysosomal membranes (Bianchi and van den Bogaart, 2020; Friedrich et al., 2012). ESCRT activity can restrict the growth of mycobacteria, presumably by mediating repair of endosomal membrane damage (Lopez-Jimenez et al., 2018; Philips et al., 2008; Portal-Celhay et al., 2016; Skowyra et al., 2018). *Mycobacterium tuberculosis* secretes the effector proteins EsxG and EsxH to prevent recruitment of host ESCRT-III proteins to phagolysosomes and promote virulence (Mehra et al., 2013; Mittal et al., 2018; Portal-Celhay et al., 2016). Similarly, *Shigella flexneri* IcsB suppresses antibacterial autophagy by interaction with the ESCRT-III protein CHMP5 (Liu et al., 2018). While the exact mechanism for ESCRT inhibition remains to be determined in these cases, they highlight the importance of pathogen engagement and modulation of the host ESCRT pathway.

Although we cannot yet say whether the potent inhibition of virus budding that we observe in cultured cells occurs *in vivo*, retroCHMP3 mRNA is expressed in squirrel monkey and mouse cell lines and well as in mouse testes and heart (**Fig. S5**). RetroCHMP3 mRNA levels appear to be low under normal conditions, but retroCHMP3^MM^ mRNA expression is modestly induced by type I interferon (**Fig. 5**), raising the possibility that retroCHMP3 levels could be regulated in specific cell types and/or during viral infection.

LINE1-mediated retrocopies are generally considered to be defective pseudogenes since the promoter and regulatory elements of parental genes are not retrocopied. Additionally, retrocopies are often truncated, either at the 5’ end due to incomplete reverse transcription during the retrotransposition event or due to premature stop codons (Casola and Betran, 2017). In some cases, however, retrocopies can acquire new expression patterns or functions (Kaessmann et al., 2009; Kubiak and Makalowska, 2017). Another ESCRT-III protein, CHMP1B, was retrotransposed in a common ancestor of placental mammals. In primates, this retrocopy replaced the parental gene and became the predominantly expressed form, demonstrating the potential of ESCRT-III retrocopies to gain regulatory elements and become functional (Ciomborowska et al., 2013). Retrotransposition events have also contributed to the evolution of other antiviral proteins: The restriction factor TRIM5α blocks replication of HIV-1 in Old World monkeys, but TRIM5α variants from New World monkeys restrict HIV-1 less efficiently (Song et al., 2005; Stremlau et al., 2004). In the New World owl monkey, however, retrotranposition of the coding sequence of the HIV-1 capsid binding factor cyclophilin A between coding exons of TRIM5α created the chimeric TRIMCyp gene that potently blocks HIV-1 replication (Nisole et al., 2004; Sayah et al., 2004). Further, some Old World macaque species encode a chimeric TRIMCyp protein that evolved by an independent retrotransposition event and restricts replication of HIV-2, feline immunodeficiency virus, and simian immunodeficiency virus SIV_AGM_tan (Brennan et al., 2008; Liao et al., 2007; Newman et al., 2008; Virgen et al., 2008; Wilson et al., 2008). Similarly, duplication of the retroviral restriction factors Fv1 in wild mice (Yap et al., 2020) and APOBEC3 in primates (Yang et al., 2020) by retrotransposition may have expanded the capacity of these proteins to restrict retroviruses by increasing the number of gene copies per genome. Retroduplication of CHMP3 in the presence of an intact parental gene appears to be another variation on this theme that allowed retroCHMP3 genes to be repurposed as virus budding inhibitors.

RetroCHMP3 proteins arose independently in at least three mammalian species by convergent evolution. We have also recently identified multiple full-length and truncated CHMP3 retrocopies in other rodent and primate genomes, including two retrocopies in mouse lemurs and dwarf lemurs that are truncated to approximately 150 amino acids (Rheinemann et al., 2021). We hypothesize that duplicated ESCRT-III proteins, and CHMP3 in particular, are especially suited to become budding inhibitors. ESCRT-III proteins consist of an N-terminal core domain that mediates filament assembly and a C-terminal autoinhibitory region. This architecture is particularly conducive to creating dominantly inhibitory ESCRT-III proteins simply by introducing a premature stop codon that removes the sites required for autoinhibition and recycling (Bajorek et al., 2009; Lata et al., 2008; Muziol et al., 2006; Shim et al., 2007; Stuchell-Brereton et al., 2007). According to our model, detoxification requires weakening of the interaction of CHMP3 with other ESCRT-III proteins. ESCRT-III proteins form highly interlocked polymers (Bertin et al., 2020; McCullough et al., 2015; Moser von Filseck et al., 2020; Nguyen et al., 2020) that can be disturbed by different combinations of amino acid substitutions along the many interaction interfaces. Therefore, attenuation of ESCRT-III filament strength can be achieved in many different ways, as evidenced by the fact that retroCHMP3 proteins in different species do not share amino acid substitutions. Retrocopies of two paralogous ESCRT-III proteins and binding partners of CHMP3, CHMP2A and CHMP4B, are much rarer in primate and rodent genomes than CHMP3 retrocopies and the retrotransposed genes that are identifiable are highly degraded in the majority of cases (Rheinemann et al., 2021). These observations indicate that CHMP3 retrocopies are either more frequently generated and/or persist longer than CHMP2A and CHMP4B retrocopies. CHMP3 is dispensable for retrovirus budding (Bartusch and Prange, 2016; Morita et al., 2011). Therefore, while the homolog of CHMP3 in yeast, Vps24, forms an essential bridge between Vps2 (CHMP2) and Snf7 (CHMP4) (Babst et al., 2002), CHMP2 and CHMP4 appear to be able to interact and form filaments in the absence of CHMP3 in mammalian cells. We speculate that this may have rendered CHMP3 particularly suitable to be repurposed as a virus budding inhibitor. However, CHMP3 depletion leads to abscission defects (Morita et al., 2010) and impairs EGFR degradation (Bache et al., 2006), and CHMP3 is therefore not dispensable for mammalian cellular ESCRT functions. Therefore, it remains unclear if the higher frequency of CHMP3 retrocopies is due to reduced CHMP3 essentiality or whether other factors such as retroduplication frequency plays a role.

In summary, we have identified retroCHMP3 proteins as broad and potent inhibitors of ESCRT-dependent virus budding. A priori, it might appear that viruses that co-opt essential cellular processes like the ESCRT pathway have found an Achilles heel that cannot easily be overcome by the host. However, retroCHMP3 proteins modify ESCRT activity to exploit a key difference in the ways that cells and viruses use the pathway. Additionally, the observation that retroCHMP3 alters ESCRT pathway function instead of targeting a viral protein raises the intriguing possibility that retroCHMP3 may be more resistant to viral counter-adaptations than other antiviral proteins that directly inhibit viral replication. Thus, retroCHMP3 proteins reveal new possibilities for blocking virus budding while maintaining cellular ESCRT functions.

## Acknowledgments

We thank Dustin Hancks for his insights and helpful discussion, Janet Iwasa for artwork, and Linda Nikolova and David Belnap at the University of Utah Electron Microscopy Core for assistance with thin-section electron microscopy. We thank Douglas Mackay and Katharine Ullman for advice and helpful discussion on cytokinesis experiments. We would also like to thank Marina Bleck for advice on live-cell imaging and Daniel Johnson and Joan Pulupa for maintenance of the TIRF microscope at Rockefeller University. Immunofluorescence microscopy was performed at the University of Utah Cell Imaging Core. Flow cytometry was performed at the University of Utah Flow Cytometry Core. This work was supported by NIH grants R37 AI 51174 (to W.I.S), P50 AI150464 (to N.C.E) and AI121880 (to A.P.S), NIH/NIGMS grant 5R01GM119585 (to S.M.S), U.S. Department of Agriculture NIFA grant 1010021 (to A.P.S.), and a Burroughs Wellcome Fund Investigators in the Pathogenesis of Infectious Disease award (to N.C.E).

## Author contributions

Conceptualization: L.R, D.M.D, G.M., W.I.S, N.C.E. Investigation: L.R, D.M.D, K.B, G.M, K.A.D, P.T, C.R.N., J.M. Supervision: A.P.S, S.M.S, W.I.S, N.C.E. Writing –Original Draft: L.R, N.C.E. Writing – Review and Editing: All authors.

## Supplemental Information

**Fig. S1:**
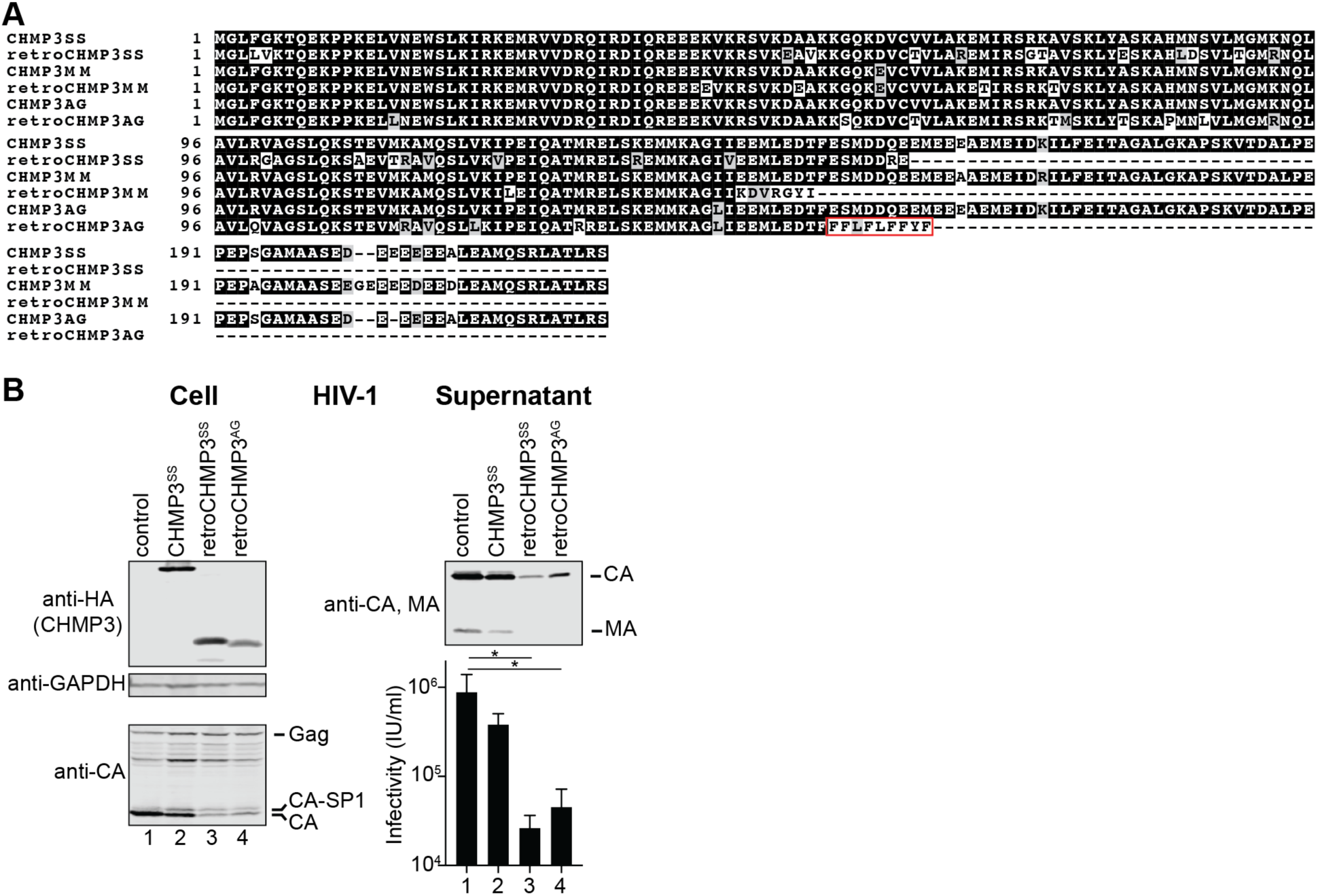
RetroCHMP3 genes in three mammalian species. Related to Fig. 1. **(A)** Alignment of parental CHMP3 and retroCHMP3 protein sequences. Note that retroCHMP3 proteins do not share amino acid substitutions. Red box in retroCHMP3^AG^ indicates sequence derived from an Alu element which was removed to enable expression. **(B)** Transient overexpression of HA-tagged retroCHMP3^AG^ inhibits release of HIV-1 proteins (CA, MA) from producer cells and reduces viral titers in supernatant. Titer graph shows mean ± SD from 3 experimental replicates. **** p<0.0001, *** p<0.001, ** p<0.01, *p<0.05 by one-way ANOVA followed by Tukey’s multiple comparisons test comparisons with p>0.05 omitted for clarity.

**Fig. S2:**
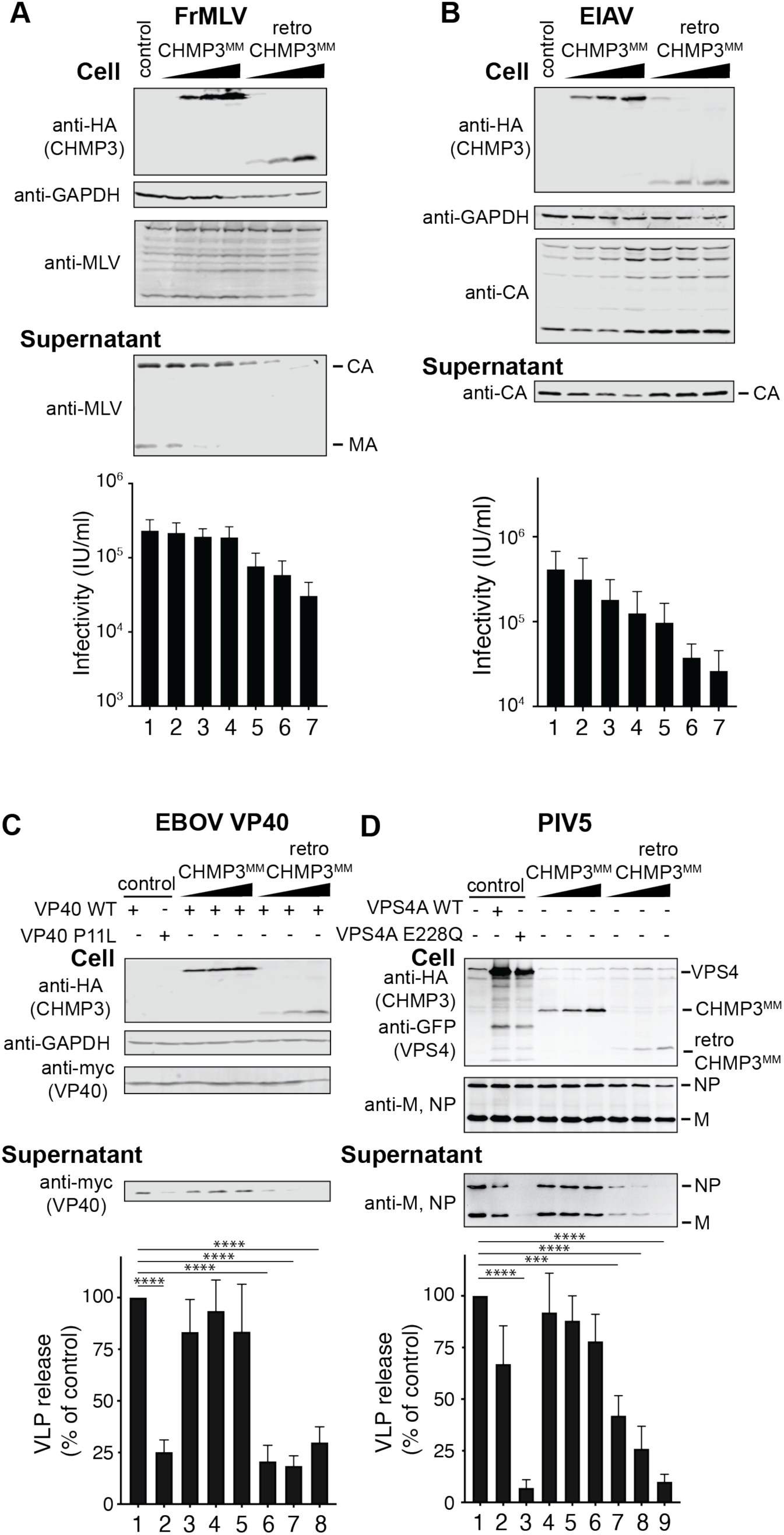
RetroCHMP3^MM^ broadly inhibits ESCRT-dependent budding. Related to Fig. 2. **(A), (B)** Transient overexpression of HA-tagged retroCHMP3^MM^ in HEK293T cells reduces the release of viral proteins and viral titers in culture supernatant of Friend murine leukemia virus (FrMLV, **A**) and equine infectious anemia virus (EIAV, **B**). Titer graphs show mean ± SD from 3 experimental replicates. **(C), (D)** Transient overexpression of HA-tagged retroCHMP3^MM^ in HEK293T cells reduces the release of Ebola virus VP40 protein **(C)** and parainfluenza virus 5 VLPs **(D)** into the supernatant. Western blots show representative results from 3 experimental replicates. Bar graphs show integrated intensities of viral protein bands in supernatant normalized to cellular fraction from 3 experimental replicates. **** p<0.0001, *** p<0.001, * p<0.05 by one-way ANOVA followed by Tukey’s multiple comparisons test. Only comparisons with the empty vector control (lane 1) are shown for clarity. For a complete set of pairwise comparisons, see Table S3.

**Fig. S3:**
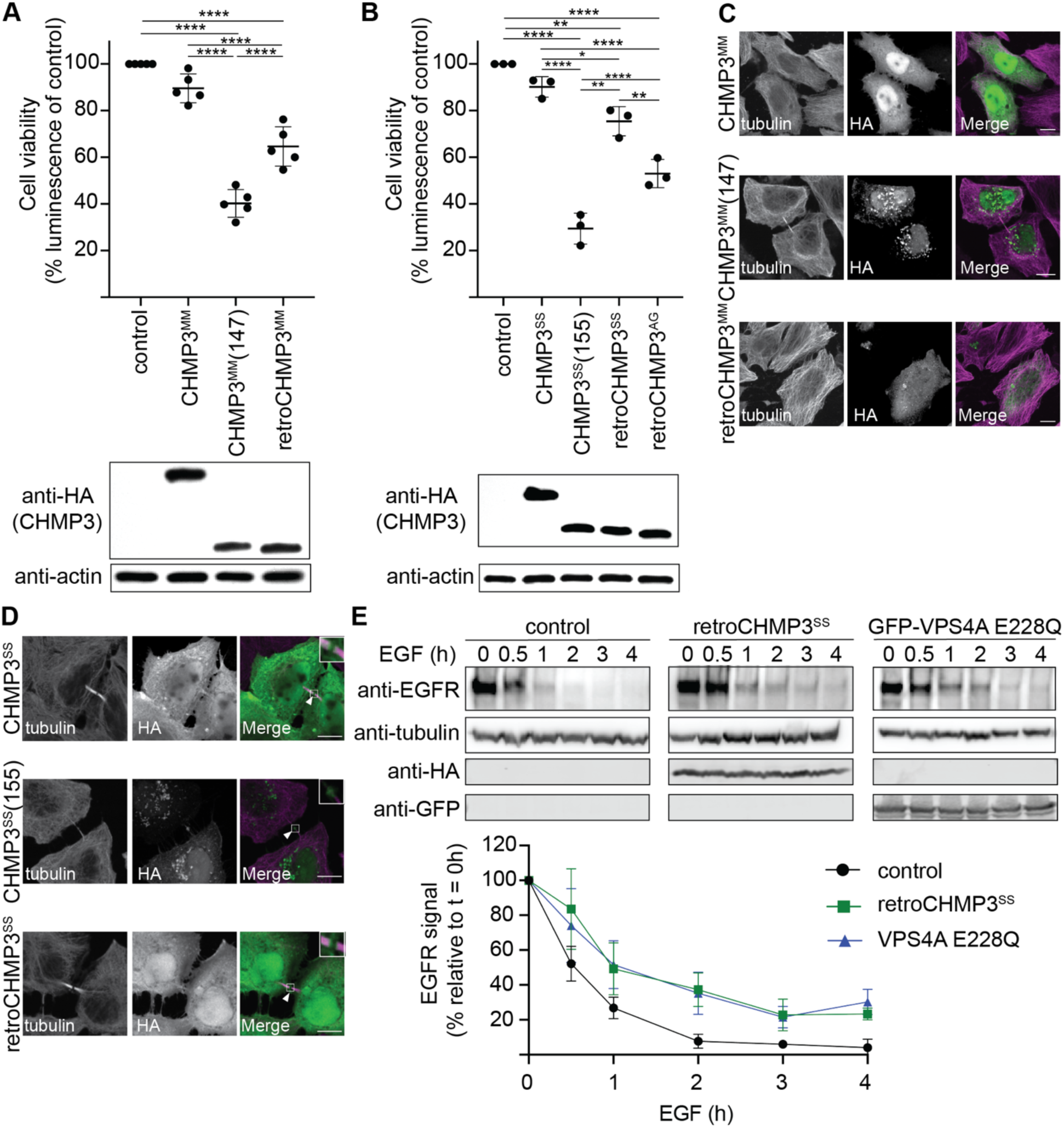
RetroCHMP3^MM^ and RetroCHMP3^AG^ evolved to lose cytotoxicity. Related to Fig. 3 **(A), (B)** Cell viability of cells transiently transfected with HA-tagged CHMP3^MM^ variants **(A)** and retroCHMP3^AG^ 24 hours after transfection. **(B)**. Mean ± SD from 5 experimental replicates with n≥4 each. **(C)** Immunofluorescence of subcellular localization of HA-tagged CHMP3^MM^ variants. Scale bar 10 μm. **(D)** Localization of HA-tagged CHMP3^SS^ variants to the midbody. White arrowhead points to CHMP3^SS^ variants at the Flemming body. Scale bar 10 μm. **(E)** Transient overexpression of HA-tagged retroCHMP3^SS^ delays EGF-induced degradation of the EGF receptor (EGFR) to a comparable degree as dominant-negative VPS4A (VPS4A-E228Q). Western blot shows representative result from 3 experimental replicates. Graph shows integrated intensities of EGFR bands normalized to intensity at t = 0h. Mean ± SD from 3 experimental replicates.

**Fig. S4:**
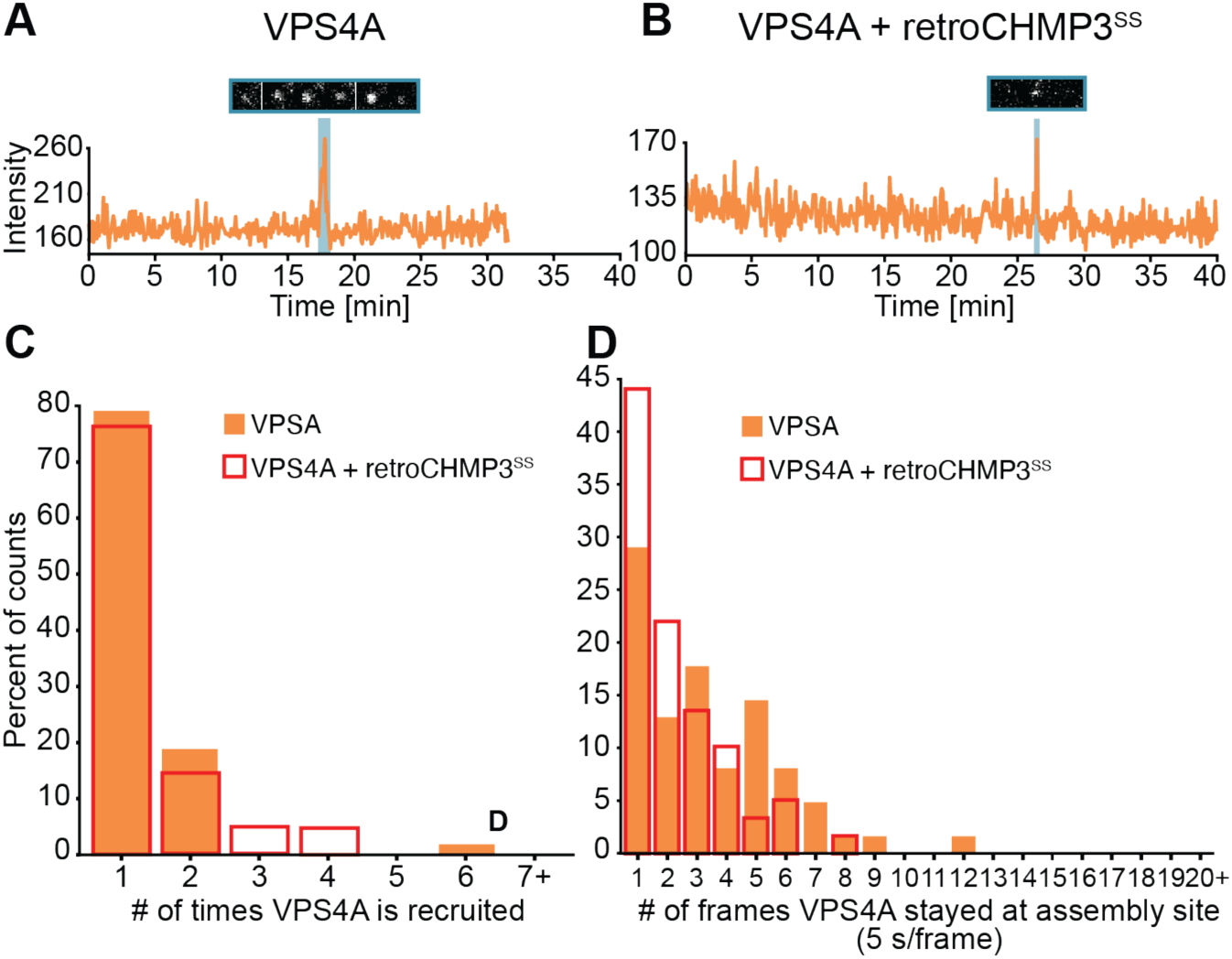
RetroCHMP3 does not affect VPS4A recruitment to HIV-1 Gag assembly sites. Related to Fig. 4. **(A), (B)** Sample trace of mCherry-VPS4A recruitment to Gag-Pol-mEGFP assembly sites in the absence **(A)** or presence **(B)** of retroCHMP3^SS^. Each numbered shaded region highlights one recruitment event and the corresponding sequence of images from that recruitment. Width of each image is 1.27 μm. **(C)** Histogram showing the number of times that VPS4A was recruited to assembly sites in the absence (n= 48 assemblies) or presence (n=42 assemblies) of retroCHMP3^SS^. p=0.349 by one-tailed T-test **(D)** Histogram showing the duration of each VPS4A recruitment in the absence (n = 62 recruitments from 48 assemblies) or presence (n = 59 recruitments from 42 assemblies) of retroCHMP3^SS^. p<0.005 by one-tailed T-test.

**Fig. S5:**
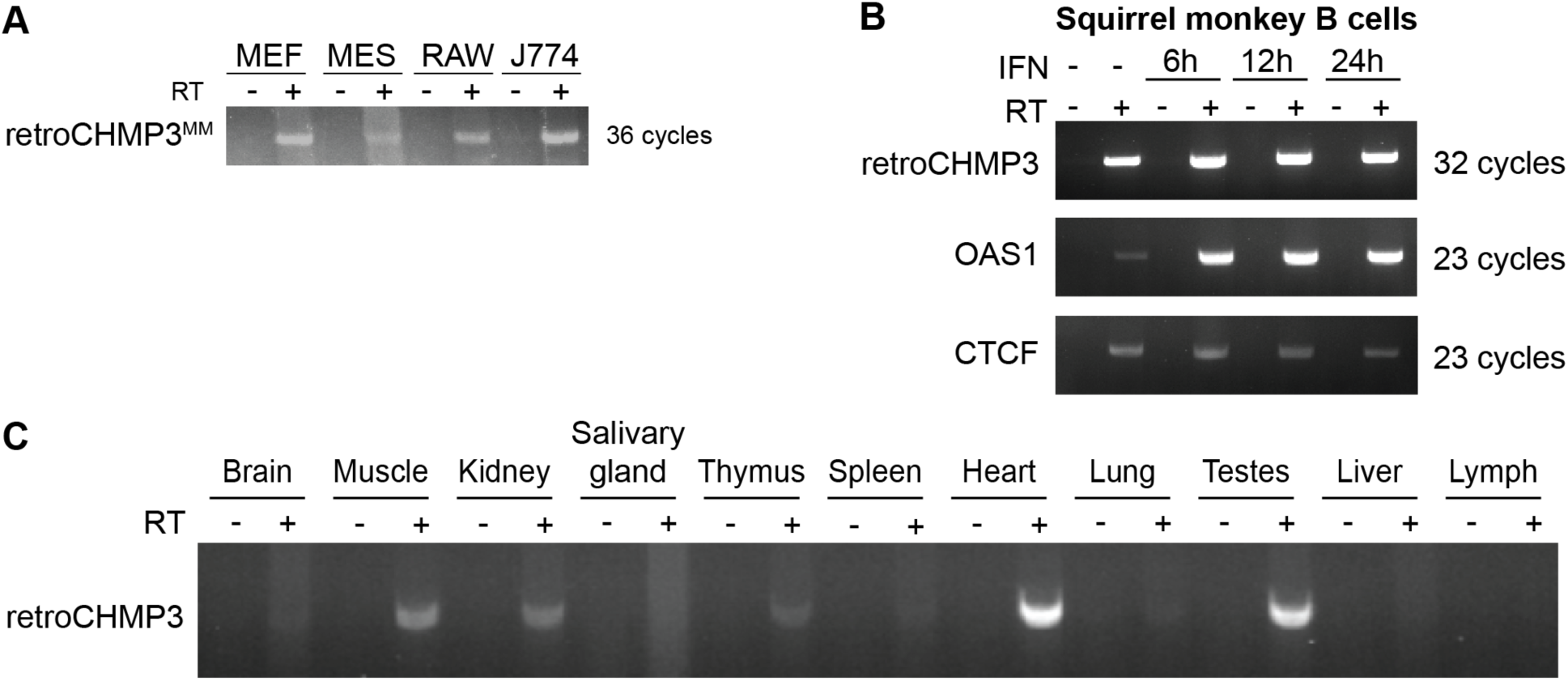
Detection of retroCHMP3 RNA. Related to Fig. 5. **(A)** Detection of retroCHMP3 RNA in mouse cell lines. MEF – mouse embryonic fibroblasts, MES – mouse embryonic stem cells, RAW – mouse macrophage cell line, J774 – mouse macrophage/monocyte cell line. **(B)** Detection of retroCHMP3 RNA in squirrel monkey B cells with and without interferon (IFN) treatment. **(C)** Detection of retroCHMP3 RNA in mouse tissues. IFN – interferon, RT – reverse transcriptase. All retroCHMP3 bands were excised, cloned and sequenced to verify a perfect match with the retroCHMP3 sequence.

**Table S1:**
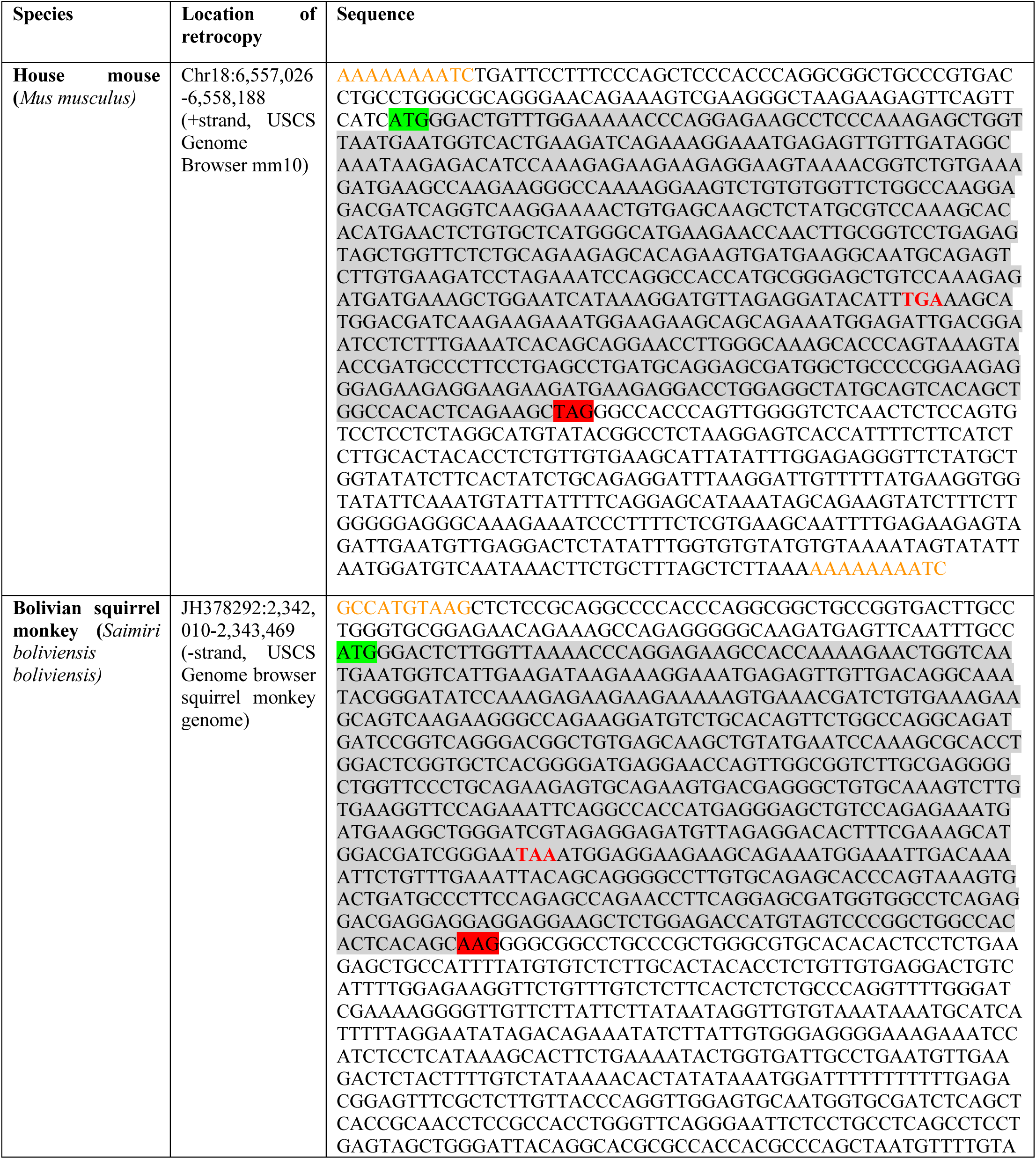
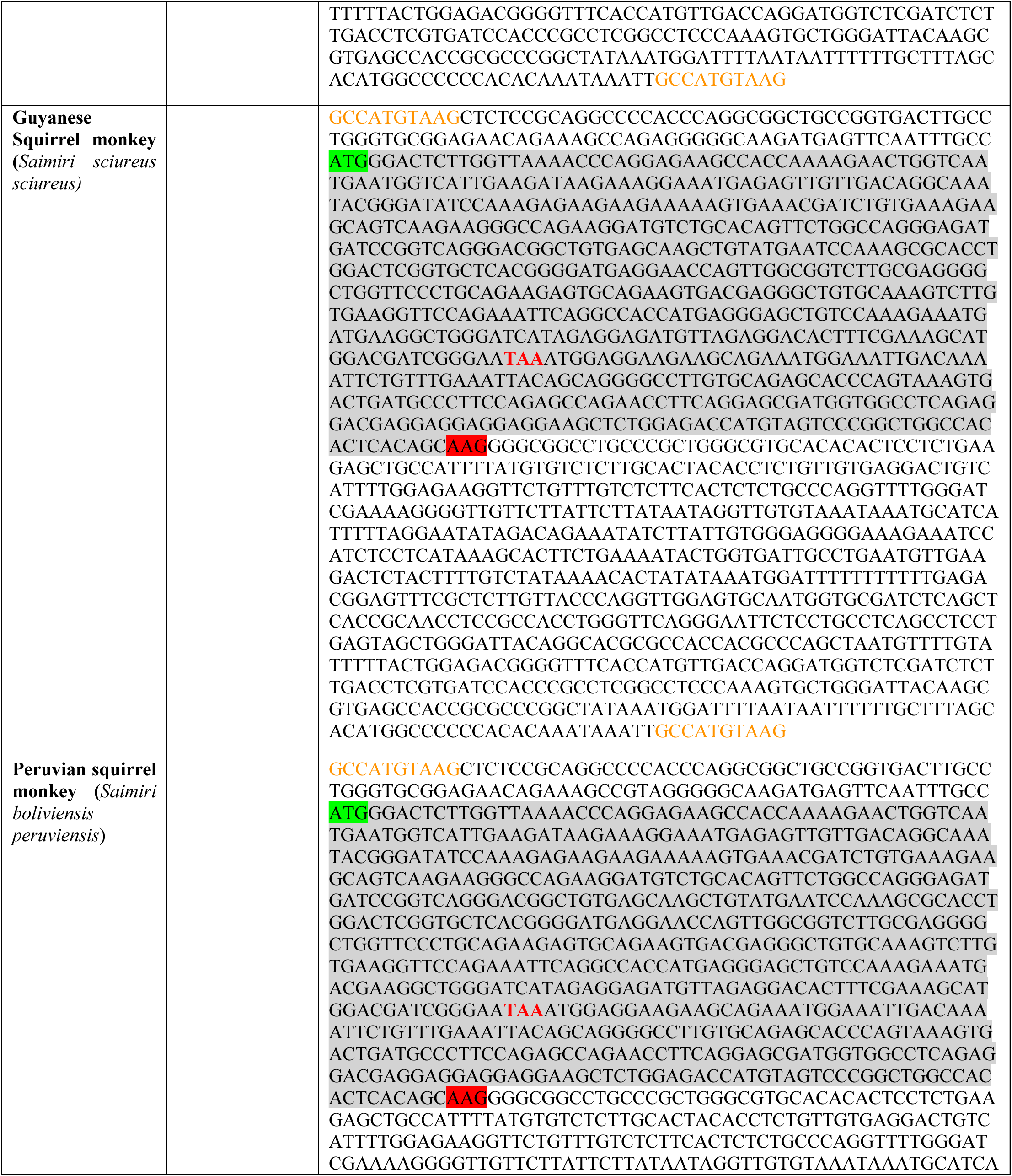
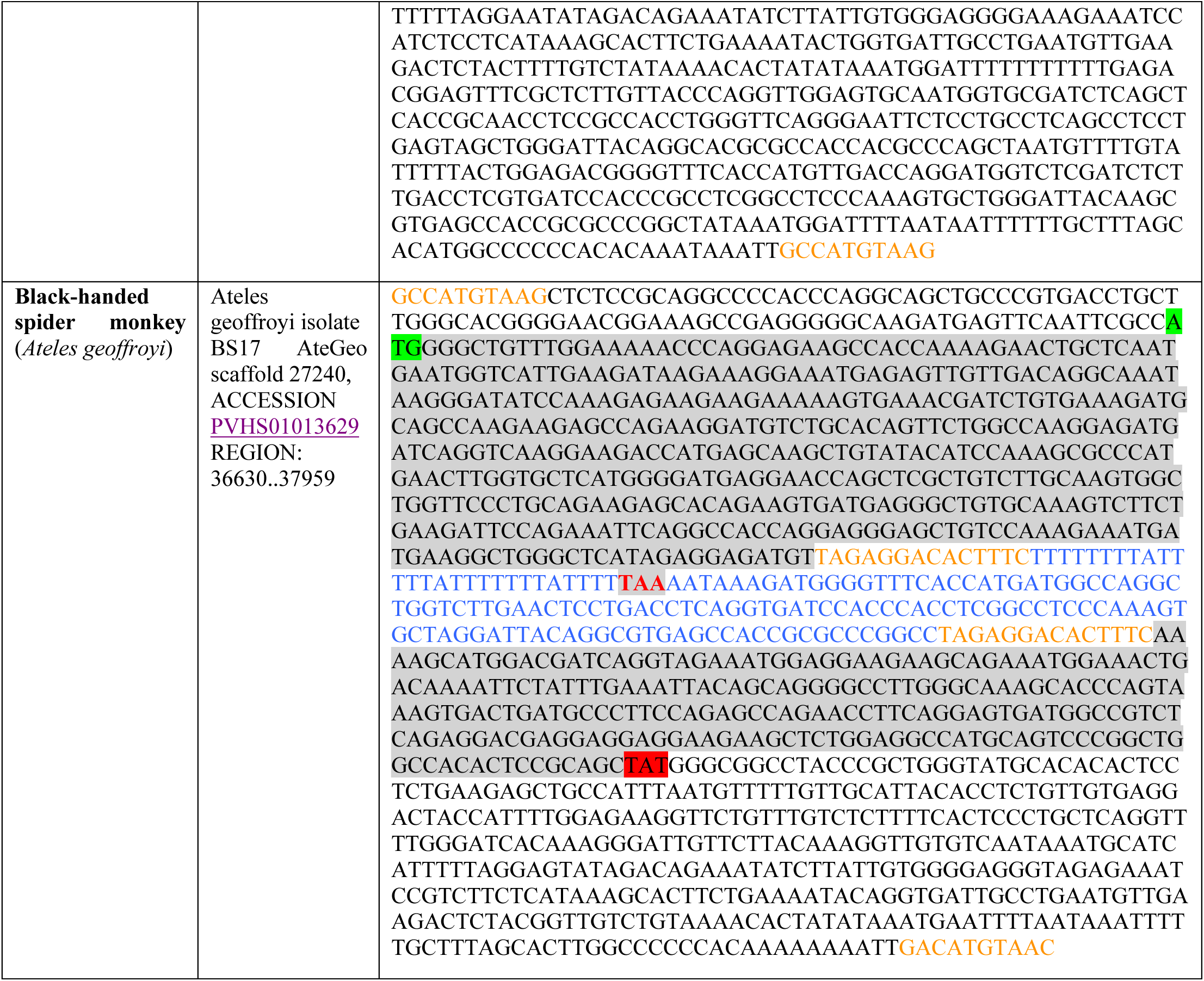
RetroCHMP3 sequences identified in this study The color coding and shading highlights key features of each retrocopy. Orange letters indicate TSD sequences flanking the retrocopy. In some cases, there is a single nucleotide difference in the 5’ and 3’ flanking TSD sequences. Gray shading indicates the original coding sequence. Green shading indicates the original start codon. Red shading indicates the location of the original stop codon. In some cases, this sequence has been modified and is therefore no longer a functional stop. Red TAG or TAA indicates location of premature, truncating stop codon. Blue letters in spider monkey sequence indicate an Alu sequence inserted within the retrocopy coding region, with orange letters indicating the TSD flanking this insertion. This Alu sequence had to be removed during cloning to enable expression.

**Table S2:**
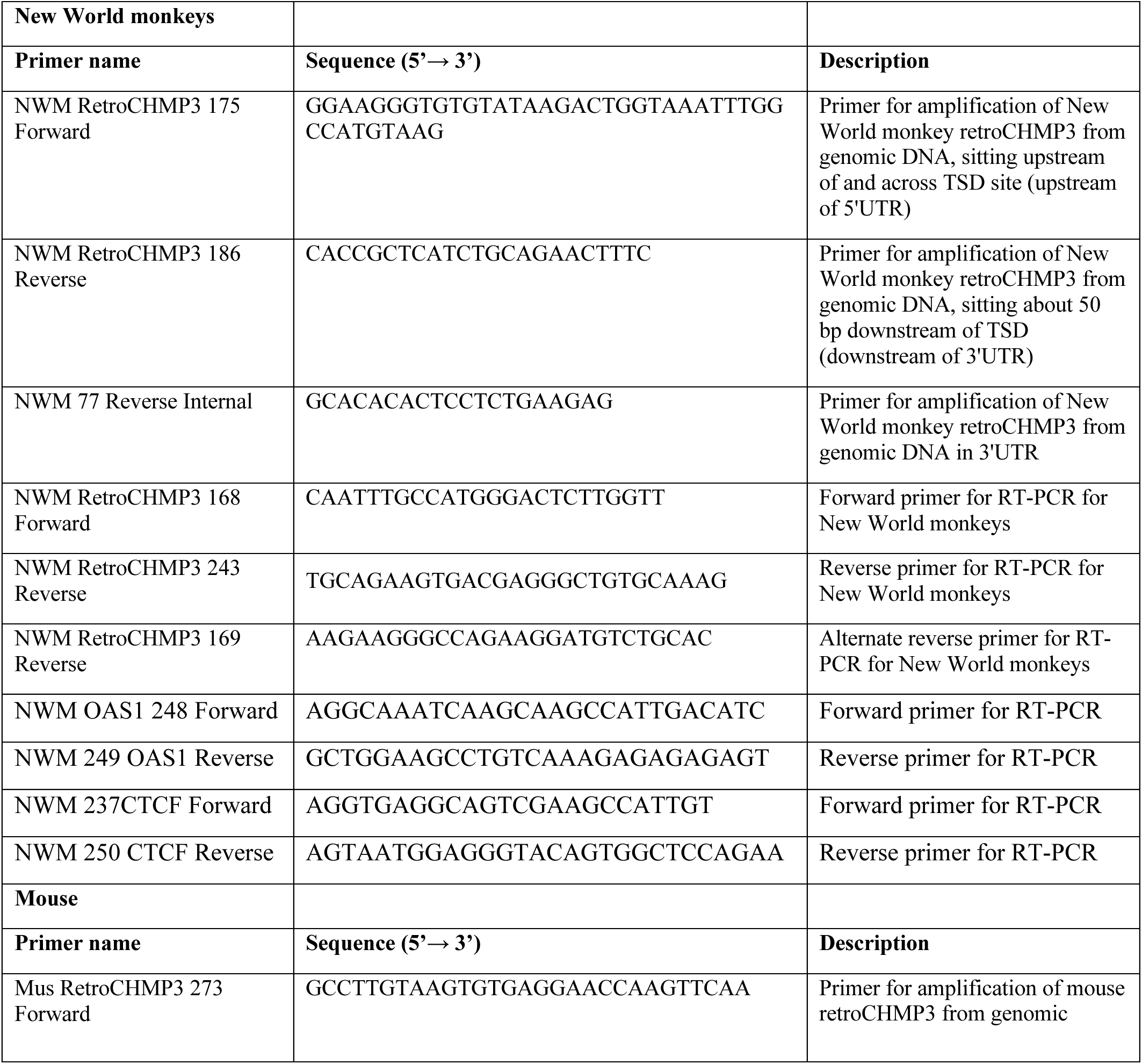
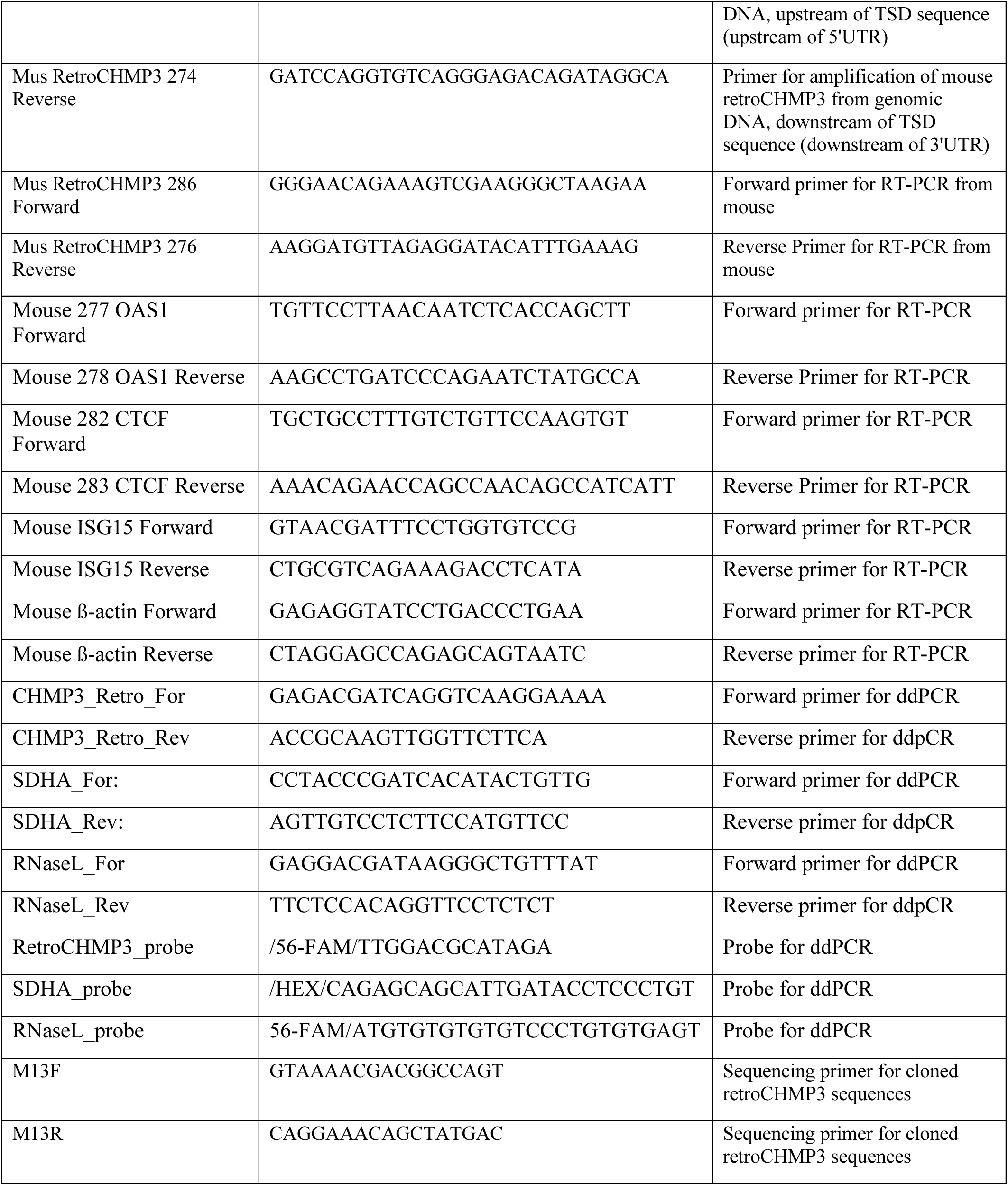
Primer sequences used in this study

**Table S3:** P-values for pairwise comparison of means for HIV-1 titers (Fig. 1C and D) and quantifications of Western Blot intensities for EBOV VP40 virus-like particles (Fig. 2C and S2C) and PIV5 virus-like particles (Fig. 2D and S2D). P-values were calculated using an ordinary one-way ANOVA followed by a Tukey’s multiple comparisons test.

| **Pairwise comparison** | **p<0.05?** | **Adjusted p-value** |
| --- | --- | --- |
| **HIV-1 + retroCHMP3<sup>SS</sup> (Fig. 1C)** |  |  |
| empty control vector (lane 1) vs. CHMP3<sup>SS</sup> 1 (lane 2) | No | 0.5628 |
| empty control vector (lane 1) vs. CHMP3<sup>SS</sup> 2 (lane 3) | No | 0.9767 |
| empty control vector (lane 1) vs. CHMP3<sup>SS</sup> 3 (lane 4) | No | 0.1191 |
| **empty control vector (lane 1) vs. retroCHMP3<sup>SS</sup> 1 (lane 5)** | **Yes** | **<0.0001** |
| **empty control vector (lane 1) vs. retro CHMP3<sup>SS</sup> 2 (lane 6)** | **Yes** | **<0.0001** |
| **empty control vector (lane 1) vs. retro CHMP3<sup>SS</sup> 3 (lane 7)** | **Yes** | **<0.0001** |
| CHMP3<sup>SS</sup> 1 (lane 2) vs. CHMP3<sup>SS</sup> 2 (lane 3) | No | 0.9517 |
| CHMP3<sup>SS</sup> 1 (lane 2) vs. CHMP3<sup>SS</sup> 3 (lane 4) | No | 0.9166 |
| **CHMP3<sup>SS</sup> 1 (lane 2) vs. retroCHMP3<sup>SS</sup> 1 (lane 5)** | **Yes** | **<0.0001** |
| **CHMP3<sup>SS</sup> 1 (lane 2) vs. retroCHMP3<sup>SS</sup> 2 (lane 6)** | **Yes** | **<0.0001** |
| **CHMP3<sup>SS</sup> 1 (lane 2) vs. retroCHMP3<sup>SS</sup> 3 (lane 7)** | **Yes** | **<0.0001** |
| CHMP3<sup>SS</sup> 2 (lane 3) vs. CHMP3<sup>SS</sup> 3 (lane 4) | No | 0.4136 |
| **CHMP3<sup>SS</sup> 2 (lane 3) vs. retroCHMP3<sup>SS</sup> 1 (lane 5)** | **Yes** | **<0.0001** |
| **CHMP3<sup>SS</sup> 2 (lane 3) vs. retroCHMP3<sup>SS</sup> 2 (lane 6)** | **Yes** | **<0.0001** |
| **CHMP3<sup>SS</sup> 2 (lane 3) vs. retroCHMP3<sup>SS</sup> 3 (lane 7)** | **Yes** | **<0.0001** |
| **CHMP3<sup>SS</sup> 3 (lane 4) vs. retroCHMP3<sup>SS</sup> 1 (lane 5)** | **Yes** | **<0.0001** |
| **CHMP3<sup>SS</sup> 3 (lane 4) vs. retroCHMP<sup>SS</sup> 2 (lane 6)** | **Yes** | **<0.0001** |
| **CHMP3<sup>SS</sup> 3 (lane 4) vs. retroCHMP3<sup>SS</sup> 3 (lane 7)** | **Yes** | **<0.0001** |
| retroCHMP3<sup>SS</sup> 1 (lane 5) vs. retroCHMP3<sup>SS</sup> 2 (lane 6) | No | >0.9999 |
| retroCHMP3<sup>SS</sup> 1 (lane 5) vs. retroCHMP3<sup>SS</sup> 3 (lane 7) | No | >0.9999 |
| retroCHMP3<sup>SS</sup> 2 (lane 6) vs. retroCHMP3<sup>SS</sup> 3 (lane 7) | No | >0.9999 |
| **HIV-1 + retroCHMP3<sup>MM</sup> (Fig. 1D)** |  |  |
| empty control vector (lane 1) vs. CHMP3<sup>MM</sup> 1 (lane 2) | No | 0.9994 |
| empty control vector (lane 1) vs. CHMP3<sup>MM</sup> 2 (lane 3) | No | 0.6855 |
| empty control vector (lane 1) vs. CHMP3<sup>MM</sup> 3 (lane 4) | No | 0.3603 |
| **empty control vector (lane 1) vs. retroCHMP3<sup>MM</sup> 1 (lane 5)** | **Yes** | **0.0002** |
| **empty control vector (lane 1) vs. retro CHMP3<sup>MM</sup> 2 (lane 6)** | **Yes** | **0.0001** |
| **empty control vector (lane 1) vs. retro CHMP3<sup>MM</sup> 3 (lane 7)** | **Yes** | **0.0001** |
| CHMP3<sup>MM</sup> 1 (lane 2) vs. CHMP3<sup>MM</sup> 2 (lane 3) | No | 0.4431 |
| CHMP3<sup>MM</sup> 1 (lane 2) vs. CHMP3<sup>MM</sup> 3 (lane 4) | No | 0.1947 |
| **CHMP3<sup>MM</sup> 1 (lane 2) vs. retroCHMP3<sup>MM</sup> 1 (lane 5)** | **Yes** | **<0.0001** |
| **CHMP3<sup>MM</sup> 1 (lane 2) vs. retroCHMP3<sup>MM</sup> 2 (lane 6)** | **Yes** | **<0.0001** |
| **CHMP3<sup>MM</sup> 1 (lane 2) vs. retroCHMP3<sup>MM</sup> 3 (lane 7)** | **Yes** | **<0.0001** |
| CHMP3<sup>MM</sup> 2 (lane 3) vs. CHMP3<sup>MM</sup> 3 (lane 4) | No | 0.9962 |
| **CHMP3<sup>MM</sup> 2 (lane 3) vs. retroCHMP3<sup>MM</sup> 1 (lane 5)** | **Yes** | **0.0024** |
| **CHMP3<sup>MM</sup> 2 (lane 3) vs. retroCHMP3<sup>MM</sup> 2 (lane 6)** | **Yes** | **0.0016** |
| **CHMP3<sup>MM</sup> 2 (lane 3) vs. retroCHMP3<sup>MM</sup> 3 (lane 7)** | **Yes** | **0.0016** |
| **CHMP3<sup>MM</sup> 3 (lane 4) vs. retroCHMP3<sup>MM</sup> 1 (lane 5)** | **Yes** | **0.0069** |
| **CHMP3<sup>MM</sup> 3 (lane 4) vs. retroCHMP3<sup>MM</sup> 2 (lane 6)** | **Yes** | **0.0045** |
| **CHMP3<sup>MM</sup> 3 (lane 4) vs. retroCHMP3<sup>MM</sup> 3 (lane 7)** | **Yes** | **0.0046** |
| retroCHMP3<sup>MM</sup> 1 (lane 5) vs. retroCHMP3<sup>MM</sup> 2 (lane 6) | No | >0.9999 |
| retroCHMP3<sup>MM</sup> 1 (lane 5) vs. retroCHMP3<sup>MM</sup> 3 (lane 7) | No | >0.9999 |
| retroCHMP3<sup>MM</sup> 2 (lane 6) vs. retroCHMP3<sup>MM</sup> 3 (lane 7) | No | >0.9999 |
| **EBOV VP40 + retroCHMP3<sup>SS</sup> (Fig. 2C)** |  |  |
| **empty control vector (lane 1) vs. VP40 P11L (lane 2)** | **Yes** | **<0.0001** |
| **empty control vector (lane 1) vs. CHMP3<sup>SS</sup> 1 (lane 3)** | **Yes** | **0.0466** |
| **empty control vector (lane 1) vs. CHMP3<sup>SS</sup> 2 (lane 4)** | **Yes** | **0.0354** |
| **empty control vector (lane 1) vs. CHMP3<sup>SS</sup> 3 (lane 5)** | **Yes** | **0.0003** |
| **empty control vector (lane 1) vs. retroCHMP3<sup>SS</sup> 1 (lane 6)** | **Yes** | **<0.0001** |
| **empty control vector (lane 1) vs. retroCHMP3<sup>SS</sup> 2 (lane 7)** | **Yes** | **<0.0001** |
| **empty control vector (lane 1) vs. retroCHMP3<sup>SS</sup> 3 (lane 8)** | **Yes** | **<0.0001** |
| **VP40 P11L (lane 2) vs. CHMP3<sup>SS</sup> 1 (lane 3)** | **Yes** | **0.0009** |
| **VP40 P11L (lane 2) vs. CHMP3<sup>SS</sup> 2 (lane 4)** | **Yes** | **0.0012** |
| VP40 P11L (lane 2) vs. CHMP3<sup>SS</sup> 3 (lane 5) | No | 0.1318 |
| VP40 P11L (lane 2) vs. retroCHMP3<sup>SS</sup> 1 (lane 6) | No | 0.8001 |
| VP40 P11L (lane 2) vs. retroCHMP3<sup>SS</sup> 2 (lane 7) | No | 0.6578 |
| VP40 P11L (lane 2) vs. retroCHMP3<sup>SS</sup> 3 (lane 8) | No | 0.9271 |
| CHMP3<sup>SS</sup> 1 (lane 3) vs. CHMP3<sup>SS</sup> 2 (lane 4) | No | >0.9999 |
| CHMP3<sup>SS</sup> 1 (lane 3) vs. CHMP3<sup>SS</sup> 3 (lane 5) | No | 0.2155 |
| **CHMP3<sup>SS</sup> 1 (lane 3) vs. retroCHMP3<sup>SS</sup> 3 (lane 6)** | **Yes** | **<0.0001** |
| **CHMP3<sup>SS</sup> 1 (lane 3) vs. retroCHMP3<sup>SS</sup> 3 (lane 7)** | **Yes** | **<0.0001** |
| **CHMP3<sup>SS</sup> 1 (lane 3) vs. retroCHMP3<sup>SS</sup> 3 (lane 8)** | **Yes** | **0.0001** |
| CHMP3<sup>SS</sup> 2 (lane 4) vs. CHMP3<sup>SS</sup> 3 (lane 5) | No | 0.2698 |
| **CHMP3<sup>SS</sup> 2 (lane 4) vs. retroCHMP3<sup>SS</sup> 1 (lane 6)** | **Yes** | **<0.0001** |
| **CHMP3<sup>SS</sup> 2 (lane 4) vs. retroCHMP3<sup>SS</sup> 2 (lane 7)** | **Yes** | **<0.0001** |
| **CHMP3<sup>SS</sup> 2 (lane 4) vs. retroCHMP3<sup>SS</sup> 3 (lane 8)** | **Yes** | **0.0001** |
| **CHMP3<sup>SS</sup> 3 (lane 5) vs. retroCHMP3<sup>SS</sup> 1 (lane 6)** | **Yes** | **0.0079** |
| **CHMP3<sup>SS</sup> 3 (lane 5) vs. retroCHMP3<sup>SS</sup> 2 (lane 7)** | **Yes** | **0.0048** |
| **CHMP3<sup>SS</sup> 3 (lane 5) vs. retroCHMP3<sup>SS</sup> 3 (lane 8)** | **Yes** | **0.0146** |
| retroCHMP3<sup>SS</sup> 1 (lane 6) vs. retroCHMP3<sup>SS</sup> 2 (lane 7) | No | >0.9999 |
| retroCHMP3<sup>SS</sup> 1 (lane 6) vs. retroCHMP3<sup>SS</sup> 3 (lane 8) | No | >0.9999 |
| retroCHMP3<sup>SS</sup> 2 (lane 7) vs. retroCHMP3<sup>SS</sup> 3 (lane 8) | No | 0.9988 |
| **EBOV VP40 + retroCHMP3<sup>MM</sup> (Fig. S2C)** |  |  |
| **empty control vector (lane 1) vs. VP40 P11L (lane 2)** | **Yes** | **<0.0001** |
| empty control vector (lane 1) vs. CHMP3<sup>SS</sup> 1 (lane 3) | No | 0.6994 |
| empty control vector (lane 1) vs. CHMP3<sup>SS</sup> 2 (lane 4) | No | 0.9972 |
| empty control vector (lane 1) vs. CHMP3<sup>SS</sup> 3 (lane 5) | No | 0.7090 |
| **empty control vector (lane 1) vs. retroCHMP3<sup>SS</sup> 1 (lane 6)** | **Yes** | **<0.0001** |
| **empty control vector (lane 1) vs. retroCHMP3<sup>SS</sup> 2 (lane 7)** | **Yes** | **<0.0001** |
| **empty control vector (lane 1) vs. retroCHMP3<sup>SS</sup> 3 (lane 8)** | **Yes** | **<0.0001** |
| **VP40 P11L (lane 2) vs. CHMP3<sup>SS</sup> 1 (lane 3)** | **Yes** | **0.0005** |
| **VP40 P11L (lane 2) vs. CHMP3<sup>SS</sup> 2 (lane 4)** | **Yes** | **<0.0001** |
| **VP40 P11L (lane 2) vs. CHMP3<sup>SS</sup> 3 (lane 5)** | **Yes** | **0.0005** |
| VP40 P11L (lane 2) vs. retroCHMP3<sup>SS</sup> 1 (lane 6) | No | 0.9997 |
| VP40 P11L (lane 2) vs. retroCHMP3<sup>SS</sup> 2 (lane 7) | No | 0.9967 |
| VP40 P11L (lane 2) vs. retroCHMP3<sup>SS</sup> 3 (lane 8) | No | 0.9996 |
| CHMP3<sup>SS</sup> 1 (lane 3) vs. CHMP3<sup>SS</sup> 2 (lane 4) | No | 0.9631 |
| CHMP3<sup>SS</sup> 1 (lane 3) vs. CHMP3<sup>SS</sup> 3 (lane 5) | No | >0.9999 |
| **CHMP3<sup>SS</sup> 1 (lane 3) vs. retroCHMP3<sup>SS</sup> 3 (lane 6)** | **Yes** | **0.0002** |
| **CHMP3<sup>SS</sup> 1 (lane 3) vs. retroCHMP3<sup>SS</sup> 3 (lane 7)** | **Yes** | **0.0001** |
| **CHMP3<sup>SS</sup> 1 (lane 3) vs. retroCHMP3<sup>SS</sup> 3 (lane 8)** | **Yes** | **0.0012** |
| CHMP3<sup>SS</sup> 2 (lane 4) vs. CHMP3<sup>SS</sup> 3 (lane 5) | No | 0.9661 |
| **CHMP3<sup>SS</sup> 2 (lane 4) vs. retroCHMP3<sup>SS</sup> 1 (lane 6)** | **Yes** | **<0.0001** |
| **CHMP3<sup>SS</sup> 2 (lane 4) vs. retroCHMP3<sup>SS</sup> 2 (lane 7)** | **Yes** | **<0.0001** |
| **CHMP3<sup>SS</sup> 2 (lane 4) vs. retroCHMP3<sup>SS</sup> 3 (lane 8)** | **Yes** | **0.0002** |
| **CHMP3<sup>SS</sup> 3 (lane 5) vs. retroCHMP3<sup>SS</sup> 1 (lane 6)** | **Yes** | **0.0002** |
| **CHMP3<sup>SS</sup> 3 (lane 5) vs. retroCHMP3<sup>SS</sup> 2 (lane 7)** | **Yes** | **0.0001** |
| **CHMP3<sup>SS</sup> 3 (lane 5) vs. retroCHMP3<sup>SS</sup> 3 (lane 8)** | **Yes** | **0.0012** |
| retroCHMP3<sup>SS</sup> 1 (lane 6) vs. retroCHMP3<sup>SS</sup> 2 (lane 7) | No | >0.9999 |
| retroCHMP3<sup>SS</sup> 1 (lane 6) vs. retroCHMP3<sup>SS</sup> 3 (lane 8) | No | 0.9787 |
| retroCHMP3<sup>SS</sup> 2 (lane 7) vs. retroCHMP3<sup>SS</sup> 3 (lane 8) | No | 0.9356 |
| **PIV5+ retroCHMP3<sup>SS</sup> (Fig. 2D)** |  |  |
| empty control vector (lane 1) vs. VPS4A WT (lane 2) | No | 0.0656 |
| **empty control vector (lane 1) vs. VPS4A E228Q (lane 3)** | **Yes** | **<0.0001** |
| empty control vector (lane 1) vs. CHMP3<sup>SS</sup> 1 (lane 4) | No | 0.2357 |
| **empty control vector (lane 1) vs. CHMP3<sup>SS</sup> 2 (lane 5)** | **Yes** | **0.0199** |
| **empty control vector (lane 1) vs. CHMP3<sup>SS</sup> 3 (lane 6)** | **Yes** | **<0.0001** |
| **empty control vector (lane 1) vs. retroCHMP3<sup>SS</sup> 1 (lane 7)** | **Yes** | **<0.0001** |
| **empty control vector (lane 1) vs. retroCHMP3<sup>SS</sup> 2 (lane 8)** | **Yes** | **<0.0001** |
| **empty control vector (lane 1) vs. retroCHMP3<sup>SS</sup> 3 (lane 9)** | **Yes** | **<0.0001** |
| **VPS4A WT (lane 2) vs. VPS4A E228Q (lane 3)** | **Yes** | **<0.0001** |
| VPS4A WT (lane 2) vs. CHMP3<sup>SS</sup> 1 (lane 4) | No | 0.9977 |
| VPS4A WT (lane 2) vs. CHMP3<sup>SS</sup> 2 (lane 5) | No | 0.9994 |
| VPS4A WT (lane 2) vs. CHMP3<sup>SS</sup> 3 (lane 6) | No | 0.0520 |
| **VPS4A WT (lane 2) vs. retroCHMP3<sup>SS</sup> 1 (lane 7)** | **Yes** | **<0.0001** |
| **VPS4A WT (lane 2) vs. retroCHMP3<sup>SS</sup> 2 (lane 8)** | **Yes** | **<0.0001** |
| **VPS4A WT (lane 2) vs. retroCHMP3<sup>SS</sup> 3 (lane 9)** | **Yes** | **<0.0001** |
| **VPS4A E228Q (lane 3) vs. CHMP3<sup>SS</sup> 1 (lane 4)** | **Yes** | **<0.0001** |
| **VPS4A E228Q (lane 3) vs. CHMP3<sup>SS</sup> 2 (lane 5)** | **Yes** | **<0.0001** |
| **VPS4A E228Q (lane 3) vs. CHMP3<sup>SS</sup> 3 (lane 6)** | **Yes** | **0.0095** |
| VPS4A E228Q (lane 3) vs. retroCHMP3<sup>SS</sup> 1 (lane 7) | No | 0.9994 |
| VPS4A E228Q (lane 3) vs. retroCHMP3<sup>SS</sup> 2 (lane 8) | No | >0.9999 |
| VPS4A E228Q (lane 3) vs. retroCHMP3<sup>SS</sup> 3 (lane 9) | No | >0.9999 |
| CHMP3<sup>SS</sup> 1 (lane 4) vs. CHMP3<sup>SS</sup> 2 (lane 5) | No | 0.9119 |
| **CHMP3<sup>SS</sup> 1 (lane 4) vs. CHMP3<sup>SS</sup> 3 (lane 6)** | **Yes** | **0.0122** |
| **CHMP3<sup>SS</sup> 1 (lane 4) vs. retroCHMP3<sup>SS</sup> 3 (lane 7)** | **Yes** | **<0.0001** |
| **CHMP3<sup>SS</sup> 1 (lane 4) vs. retroCHMP3<sup>SS</sup> 3 (lane 8)** | **Yes** | **<0.0001** |
| **CHMP3<sup>SS</sup> 1 (lane 4) vs. retroCHMP3<sup>SS</sup> 3 (lane 9)** | **Yes** | **<0.0001** |
| CHMP3<sup>SS</sup> 2 (lane 5) vs. CHMP3<sup>SS</sup> 3 (lane 6) | No | 0.1581 |
| **CHMP3<sup>SS</sup> 2 (lane 5) vs. retroCHMP3<sup>SS</sup> 1 (lane 7)** | **Yes** | **<0.0001** |
| **CHMP3<sup>SS</sup> 2 (lane 5) vs. retroCHMP3<sup>SS</sup> 2 (lane 8)** | **Yes** | **<0.0001** |
| **CHMP3<sup>SS</sup> 2 (lane 5) vs. retroCHMP3<sup>SS</sup> 3 (lane 9)** | **Yes** | **<0.0001** |
| **CHMP3<sup>SS</sup> 3 (lane 6) vs. retroCHMP3<sup>SS</sup> 1 (lane 7)** | **Yes** | **0.0324** |
| **CHMP3<sup>SS</sup> 3 (lane 6) vs. retroCHMP3<sup>SS</sup> 2 (lane 8)** | **Yes** | **0.0122** |
| **CHMP3<sup>SS</sup> 3 (lane 6) vs. retroCHMP3<sup>SS</sup> 3 (lane 9)** | **Yes** | **0.0095** |
| retroCHMP3<sup>SS</sup> 1 (lane 7) vs. retroCHMP3<sup>SS</sup> 2 (lane 8) | No | 0.9999 |
| retroCHMP3<sup>SS</sup> 1 (lane 7) vs. retroCHMP3<sup>SS</sup> 3 (lane 9) | No | 0.9994 |
| retroCHMP3<sup>SS</sup> 2 (lane 8) vs. retroCHMP3<sup>SS</sup> 3 (lane 9) | No | >0.9999 |
| **PIV5+ retroCHMP3<sup>MM</sup> (Fig. S2D)** |  |  |
| empty control vector (lane 1) vs. VPS4A WT (lane 2) | No | 0.0594 |
| **empty control vector (lane 1) vs. VPS4A E228Q (lane 3)** | **Yes** | **<0.0001** |
| empty control vector (lane 1) vs. CHMP3<sup>SS</sup> 1 (lane 4) | No | 0.9942 |
| empty control vector (lane 1) vs. CHMP3<sup>SS</sup> 2 (lane 5) | No | 0.9357 |
| empty control vector (lane 1) vs. CHMP3<sup>SS</sup> 3 (lane 6) | No | 0.4030 |
| **empty control vector (lane 1) vs. retroCHMP3<sup>SS</sup> 1 (lane 7)** | **Yes** | **0.0003** |
| **empty control vector (lane 1) vs. retroCHMP3<sup>SS</sup> 2 (lane 8)** | **Yes** | **<0.0001** |
| **empty control vector (lane 1) vs. retroCHMP3<sup>SS</sup> 3 (lane 9)** | **Yes** | **<0.0001** |
| **VPS4A WT (lane 2) vs. VPS4A E228Q (lane 3)** | **Yes** | **0.0002** |
| VPS4A WT (lane 2) vs. CHMP3<sup>SS</sup> 1 (lane 4) | No | 0.2573 |
| VPS4A WT (lane 2) vs. CHMP3<sup>SS</sup> 2 (lane 5) | No | 0.4600 |
| VPS4A WT (lane 2) vs. CHMP3<sup>SS</sup> 3 (lane 6) | No | 0.9597 |
| VPS4A WT (lane 2) vs. retroCHMP3<sup>SS</sup> 1 (lane 7) | No | 0.2573 |
| **VPS4A WT (lane 2) vs. retroCHMP3<sup>SS</sup> 2 (lane 8)** | **Yes** | **0.0112** |
| **VPS4A WT (lane 2) vs. retroCHMP3<sup>SS</sup> 3 (lane 9)** | **Yes** | **0.0004** |
| **VPS4A E228Q (lane 3) vs. CHMP3<sup>SS</sup> 1 (lane 4)** | **Yes** | **<0.0001** |
| **VPS4A E228Q (lane 3) vs. CHMP3<sup>SS</sup> 2 (lane 5)** | **Yes** | **<0.0001** |
| **VPS4A E228Q (lane 3) vs. CHMP3<sup>SS</sup> 3 (lane 6)** | **Yes** | **<0.0001** |
| **VPS4A E228Q (lane 3) vs. retroCHMP3<sup>SS</sup> 1 (lane 7)** | **Yes** | **0.0396** |
| VPS4A E228Q (lane 3) vs. retroCHMP3<sup>SS</sup> 2 (lane 8) | No | 0.5822 |
| VPS4A E228Q (lane 3) vs. retroCHMP3<sup>SS</sup> 3 (lane 9) | No | >0.9999 |
| CHMP3<sup>SS</sup> 1 (lane 4) vs. CHMP3<sup>SS</sup> 2 (lane 5) | No | >0.9999 |
| CHMP3<sup>SS</sup> 1 (lane 4) vs. CHMP3<sup>SS</sup> 3 (lane 6) | No | 0.8642 |
| **CHMP3<sup>SS</sup> 1 (lane 4) vs. retroCHMP3<sup>SS</sup> 3 (lane 7)** | **Yes** | **0.0016** |
| **CHMP3<sup>SS</sup> 1 (lane 4) vs. retroCHMP3<sup>SS</sup> 3 (lane 8)** | **Yes** | **<0.0001** |
| **CHMP3<sup>SS</sup> 1 (lane 4) vs. retroCHMP3<sup>SS</sup> 3 (lane 9)** | **Yes** | **<0.0001** |
| CHMP3<sup>SS</sup> 2 (lane 5) vs. CHMP3<sup>SS</sup> 3 (lane 6) | No | 0.9767 |
| **CHMP3<sup>SS</sup> 2 (lane 5) vs. retroCHMP3<sup>SS</sup> 1 (lane 7)** | **Yes** | **0.0039** |
| **CHMP3<sup>SS</sup> 2 (lane 5) vs. retroCHMP3<sup>SS</sup> 2 (lane 8)** | **Yes** | **0.0001** |
| **CHMP3<sup>SS</sup> 2 (lane 5) vs. retroCHMP3<sup>SS</sup> 3 (lane 9)** | **Yes** | **<0.0001** |
| **CHMP3<sup>SS</sup> 3 (lane 6) vs. retroCHMP3<sup>SS</sup> 1 (lane 7)** | **Yes** | **0.0323** |
| **CHMP3<sup>SS</sup> 3 (lane 6) vs. retroCHMP3<sup>SS</sup> 2 (lane 8)** | **Yes** | **0.0011** |
| **CHMP3<sup>SS</sup> 3 (lane 6) vs. retroCHMP3<sup>SS</sup> 3 (lane 9)** | **Yes** | **<0.0001** |
| retroCHMP3<sup>SS</sup> 1 (lane 7) vs. retroCHMP3<sup>SS</sup> 2 (lane 8) | No | 0.7638 |
| retroCHMP3<sup>SS</sup> 1 (lane 7) vs. retroCHMP3<sup>SS</sup> 3 (lane 9) | No | 0.0725 |
| retroCHMP3<sup>SS</sup> 2 (lane 8) vs. retroCHMP3<sup>SS</sup> 3 (lane 9) | No | 0.7638 |

## STAR Methods

### RESOURCE AVAILABILITY

#### Lead Contact

Further information and request for resources and reagents should be directed to and will be fulfilled by the Lead Contact, Nels C. Elde.

#### Materials Availability Statement

Plasmids generated in this study have been deposited to Addgene. The published article includes all sequencing data generated during this study.

#### Data and Code Availability

This study did not generate or analyze datasets or code.

### EXPERIMENTAL MODEL AND SUBJECT DETAILS

#### Cells

HEK293T cells (CRL-3216, Human embryonic kidney endothelial cells, sex: female), GSML cells (CRL-2699, squirrel monkey B lymphoblast cell line, sex: male), SML cells (CRL-2311, squirrel monkey lymphocytes cell line, clone 4D8, sex: male), RAW 264.7 cells (TIB-71, mouse macrophage cell line, sex: male) and J774A.1 cells (TIB-67, mouse monocyte/macrophage cells, sex: female) were obtained from ATCC. MCEC cells (mouse cardiac endothelial cells, CLU510, sex: unknown) were obtained from Cedarlane. MT-4 cells (Human T cells, NIH-ARP 120, sex: male) were obtained from the NIH AIDS Reagent program. HeLa N cells (human cervical epithelial cells, sex: female) were a kind gift from Katharine Ullman (University of Utah). DFJ8 cells (DF-1 chicken embryonic fibroblasts stably expressing MCAT-1, sex: unknown) were a kind gift from Walther Mothes (Yale University). MES cells (mouse embryonic stem cells, line G4-56, passage 14, sex: unknown) were a kind gift from Mario Capecchi (University of Utah). MEF cells (mouse embryonic fibroblasts, sex: unknown) were isolated from WT E13.5 C57BL/6 embryos and immortalized by lentiviral transduction with large T antigen (Zhu et al., 1991).

HEK293T, DFJ8, HeLa N, MCEC and MEFs were maintained in Dulbecco’s modified Eagle’s medium (DMEM, Gibco, Thermo Fisher Scientific) containing 10% FBS at 37°C and 5% CO_2_. RAW 264.7 cells and J774A.1 cells were maintained in DMEM (Gibco, Thermo Fisher Scientific) containing 5% FBS at 37°C and 5% CO_2_. MT-4 and squirrel monkey B lymphoblast cells were maintained in RPMI-1640 medium (Gibco, Thermo Fisher Scientific) containing 10% FBS (Atlanta Biologicals) at 37°C and 5% CO_2_. MES cells were maintained as previously described (George et al., 2007).

Cells were tested for mycoplasma contamination every 3 months using the PCR Mycoplasma Detection Kit (abm). Cells have not been authenticated.

#### Genomic DNA sources

Sources for genomic DNA were as follows: Blood samples from *Cebus apella* (tufted capuchin, three individuals) were obtained from the NIH Animal Center (Dickerson MD, USA). Blood samples from *Ateles geoffroyi* (black-handed spider monkey, one individual) and *Ateles fusciceps* (brown-headed spider monkey, one individual) were obtained from Utah’s Hogle Zoo (Salt Lake City, UT, USA). Blood samples from *Saimiri boliviensis boliviensis* (Bolivian squirrel monkey, 6 individuals), *Saimiri boliviensis peruviensis* (Peruvian squirrel monkey, 6 individuals) and *Saimiri sciureus sciureus* (Guyanese squirrel monkey, 6 individuals) were obtained from the Institute Pasteur de la Guyane (Cayenne Cedex, French Guiana). Primary fibroblast cell lines and genomic DNA from *Saimiri sciureus* (common squirrel monkey, AG05311) were obtained from Coriell Cell Repositories. Blood samples from *Mus musculus* (house mouse, C57BL/6) were obtained from the University of Utah animal facility (Salt Lake City, UT, USA).

### METHOD DETAILS

#### Identification, sequencing and cloning of CHMP3 retrocopies

Retrocopies of CHMP3 were identified using BLAT on the UCSC genome browser (http://genome.ucsc.edu/index.html) or NCBI BLAST (https://blast.ncbi.nlm.nih.gov/Blast.cgi). Specifically, the full-length CHMP3 cDNA (5’ UTR, coding sequence and 3’UTR) encoded by *Homo sapiens* (ENST00000263856.9, https://www.ncbi.nlm.nih.gov/nuccore/NM_016079.4) or *Mus musculus* (ENST00000263856.9, https://www.ncbi.nlm.nih.gov/nuccore/NM_025783.4) were used as queries against mammalian genomes on the UCSC browser or whole genome shotgun contigs via NCBI BLAST to retrieve retrogenes present in the corresponding genomes. Candidate retrocopies were identified as CHMP3 sequences lacking introns – a key structural characteristic of processed pseudogenes. In all cases, we have identified the target site duplication flanking each retrocopy.

Retrogene insertions in New World monkeys and Mus genomes were validated by PCR amplification of genomic DNA using primers positioned in flanking sequences upstream and downstream of the target-site duplication. For those species where no genomic DNA was available, sequences are from publicly available whole genome shotgun contigs on NCBI BLAST. Primer sequences used are listed in Table S2. PCR products were resolved on 2% agarose gels followed by extraction and purification of candidate DNA amplicons using Zymoclean Gel DNA Recovery Kits (Zymo Research). Purified PCR products were cloned using the TOPO TA cloning kit (Invitrogen), following the manufacturer’s instructions. Following transformation and plating, bacterial colonies were selected and grown in overnight cultures for plasmid purification using the Zyppy Plasmid Miniprep kit (Zymo Research). Sanger sequencing of clones was performed using M13F and M13R primers. RetroCHMP3 sequences are reported in Table S1.

#### Plasmids

The expression constructs used in this study are described in the Key Resources Table. All newly generated expression constructs have been submitted to the Addgene plasmid repository (https://www.addgene.org/). The HIV-1 expression construct NLENG1-IRES-GFP was a kind gift from David N. Levy (New York University) (Levy et al., 2004). The FrMLV expression vector pLRB303 FrMLV PRR Env(273)GFP was a kind gift from Walther Mothes (Yale University) (Lehmann et al., 2005; Sherer et al., 2003). The pLNCX2-mCherry-CHMP4B and pLNCX2-mCherry-VPS4A plasmids were previously described (Bleck et al., 2014; Johnson et al., 2018). pCRV1-NL4.3GagPol-mEGFP and GagPol-mEGFP-D25A were created from the pCRV1-NL4.3-GagPol plasmid, which was a kind gift from Paul Bieniasz (Rockefeller University), by inserting mEGFP in the matrix domain with a matrix/capsid cleavage site on either side of the mEGFP.

#### HIV-1 budding assays

0.8 × 10^6^ HEK293T cells were seeded in 6-well plates 18-24 h before transfection. Cells were co-transfected with 1 μg HIV-1 NLENG1-IRES-GFP expression vector and increasing amounts of expression vectors for full-length CHMP3 or retroCHMP3 to generate an expression gradient using 10 μl Lipofectamine 2000 (Thermo Fisher Scientific) according to manufacturer’s instructions. Plasmid DNA amounts were as follows: 2 μg pCMV(Δ3)-squirrel-monkey-full-length-CHMP3-HA, 2 μg pCMV(Δ2)-squirrel-monkey-full-length-CHMP3-HA, 2 μg pCMV(Δ1)-squirrel-monkey-full-length CHMP3-HA; 500 ng, 1 μg and 1.5 μg pEF1α-squirrel-monkey-retroCHMP3-HA; 100 ng, 250 ng and 500 ng pCMV(WT)-mouse-full-length-CHMP3-HA; 500 ng, 1 μg and 1.5 μg pEF1α-mouse-retroCHMP3-HA, 2 μg pEF1α-spider-monkey-retroCHMP3-HA. If necessary, plasmid amounts were adjusted with empty pCMV(WT) vector. The medium was replaced with 2 ml DMEM 4-6 h later. Cells were harvested for western blot analysis and supernatants were harvested for titer measurements and western blot analysis as described below 24 h post-transfection.

For western blot analyses, virions from 1 ml supernatant were pelleted by centrifugation through a 200 μl 20% sucrose cushion (90 min, 15,000×*g*, 4°C) and denatured by adding 50 μl 1x Laemmli SDS-PAGE loading buffer and boiling for 5 min. Cells were washed in 1 ml PBS, lysed for 5min on ice in 200 μl Triton lysis buffer (50 mM Tris pH 7.4, 150 mM NaCl, 1% Triton X-100) supplemented with mammalian protease inhibitor (Sigma-Aldrich). 150 μl 2× Laemmli SDS-PAGE loading buffer supplemented with 10% 2-mercaptoethanol (Sigma-Aldrich) were added, and samples were boiled for 10 min. Proteins were separated by SDS-PAGE, transferred onto PVDF membranes and probed with antibodies. Primary antibodies were as follows: anti-HIV CA (Covance, UT415), anti-HIV MA (Covance, UT556), anti-HA (Sigma-Aldrich, H6908), anti-GAPDH (EMD Millipore, MAB374). Antibodies with UT numbers were raised against recombinant proteins purified in the Sundquist laboratory. Bands were visualized by probing the membrane with fluorescently labeled secondary antibodies (Li-Cor Biosciences) and scanning with an Odyssey Imager (Li-Cor Biosciences).

HIV-1 titers were assayed in MT-4 cells by quantifying GFP expression in target cells. Briefly, MT-4 cells (100,000 cells/well, 96-well plate) were infected with three different dilutions of virus-containing culture media in duplicate. After 16 h, dextran sulfate was added to a final concentration of 100 μg/ml. Cells were harvested after 48 h, washed in PBS three times and fixed with 2% paraformaldehyde for 20 min at room temperature. Percentages of GFP-positive cells were determined by flow cytometry on a FACSCanto (BD Biosciences).

#### FrMLV, EIAV and EBOV Vp40 budding assays

5 × 10^5^ HEK293T cells (FrMLV, EIAV) or 8 × 10^5^ HEK293T cells (EBOV Vp40) were seeded in 6-well plates 18-24 h before transfection. Cells were co-transfected with 2 μg FrMLV-GFP expression vector pLRB303 FrMLV PRR Env(273)GFP (Lehmann et al., 2005; Sherer et al., 2003) or an EIAV vector system (Olsen, 1998; Yee et al., 1994) comprising 0.2 μg pEV53, 0.2 μg pSIN6.1CeGFPW and 0.075 μg phCMV-VSV-G, or 1 μg Myc-EbVp40-WT (Martin-Serrano et al., 2001) or Myc-EbVp40-P11L (Martin-Serrano et al., 2004) expression vector and increasing amounts of expression vectors for full-length CHMP3 or retroCHMP3 to generate an expression gradient using 10 μl Lipofectamine 2000 (Thermo Fisher Scientific) according to manufacturer’s instructions. Plasmid DNA amounts were as follows: 2 μg pCMV(Δ3)-squirrel-monkey-full-length-CHMP3-HA, 2 μg pCMV(Δ2)-squirrel-monkey-full-length-CHMP3-HA, 2 μg pCMV(Δ1)-squirrel-monkey-full-length CHMP3-HA; 500 ng, 1 μg and 1.5 μg pEF1α-squirrel-monkey-retroCHMP3-HA; 100 ng, 250 ng and 500 ng pCMV(WT)-mouse-full-length-CHMP3-HA; 500 ng, 1 μg and 1.5 μg pEF1α-mouse-retroCHMP3-HA. If necessary, plasmid amounts were adjusted with empty pCMV(WT) vector. The medium was replaced with 2 ml DMEM 4-6 h later. Cells were harvested for western blot analysis and supernatants were harvested for titer measurements and western blot analysis as described below 48 h (EIAV and FrMLV) or 18 h (EbVp40) post-transfection.

Western blot analyses were carried out as described for HIV-1. Primary antibodies were as follows: anti-EIAV CA (UT418), anti-FriendMLV (ATCC, VR-1537), anti-HA (Sigma-Aldrich, H6908), anti-myc (EMD Millipore, 05-724), anti-GAPDH (EMD Millipore, MAB374). Bands were visualized by probing the membrane with fluorescently labeled secondary antibodies (Li-Cor Biosciences) and scanning with an Odyssey Imager (LI-COR Biosciences).

Titers for the EIAV vector and FrMLV were assayed in HEK293T or DFJ cells, respectively, by quantifying GFP expression in target cells. Briefly, cells (5 × 10^4^ cells/well, 24-well plate) were infected with three different dilutions of virus-containing culture media. For FrMLV, 4 μg/ml polybrene was added during infection. After 16 h, fresh growth medium was added. Cells were harvested after 48 h using 0.25% trypsin-EDTA (Gibco, ThermoFisher Scientific) and fixed with 2% paraformaldehyde for 20 min at RT. Percentages of GFP-positive cells were determined by flow cytometry on a FACSCanto (BD Biosciences).

For EBOV Vp40 release, band intensities from Western Blots of supernatants were quantified using LI-COR Image Studio software and normalized to band intensities of cellular fractions.

#### PIV-5 budding assays

For PIV5 budding assays, HEK293T cells in 6-well plates were transfected at 70-80% confluency using Lipofectamine-Plus reagents (Thermo Fisher Scientific) according to manufacturer’s instructions. Cells were transfected with plasmids encoding PIV5 M, NP, and HN proteins for generation of virus-like particles as described previously (Schmitt et al., 2002), together with plasmids encoding full-length CHMP3, retroCHMP3, VPS4, or VPS4-E228Q. Plasmid amounts were as follows: 300 ng pCAGGS-PIV5 M, 100 ng pCAGGS-PIV5 NP, 750 ng pCAGGS-PIV5 HN, 100 ng pGFP-VPS4A, 100 ng pGFP-VPS4A E228Q, 2 μg pCMV(Δ3)-squirrel-monkey-full-length-CHMP3-HA, 2 μg pCMV(Δ2)-squirrel-monkey full-length-CHMP3-HA, 2 μg pCMV(Δ1)-squirrel-monkey-full-length-CHMP3-HA; 500 ng, 1 μg and 1.5 μg pEF1α-squirrel-monkey-retroCHMP3-HA. Total plasmid quantities were equalized using pCMV empty vector. After overnight incubation, the culture medium was replaced with DMEM supplemented with 2% fetal bovine serum. 40 h post-transfection, cell and culture media fractions were collected. Cells were pelleted and lysed in SDS-PAGE loading buffer containing 2.5% (wt/vol) DTT. Virus-like particles from culture media fractions were pelleted through 20% sucrose cushions, resuspended, floated to the tops of sucrose flotation gradients, pelleted again, and then resuspended in SDS-PAGE loading buffer containing 2.5% (wt/vol) DTT. Proteins from cell lysates and virus-like particles were fractionated using 10% or 15% SDS-PAGE and transferred to PVDF membranes for immunodetection. Primary antibodies were as follows: anti-PIV5 M (M-f) (Randall et al., 1987), anti-PIV5 NP (NP-a) (Randall et al., 1987), anti-HA (Proteintech 51064-2-AP), anti-GFP (Thermo Fisher Scientific, A-6455). Bands were visualized by probing with alkaline phosphatase-conjugated secondary antibodies followed by incubation with Vistra ECF substrate (GE Healthcare) and imaging using a Fuji FLA-7000 phosphorimager. VLP production efficiency was calculated as the quantity of M protein in purified VLPs divided by the quantity of M protein in the corresponding cell lysate fraction, normalized to the value obtained in the absence of CHMP3 or VPS4A expression.

#### Transmission electron microscopy (TEM)

For thin sectioning, 0.8 × 10^6^ HEK293T cells were seeded in 6-well plates 18-24 h before transfection. Cells were co-transfected with the designated expression plasmids as described above. 24 h post-transfection, the media was removed and cells were harvested in 1 ml fixation buffer (2.5% glutaraldehyde/1% paraformaldehyde in sodium cacodylate buffer [50 mM sodium cacodylate, 18 mM sucrose, 2 mM CaCl_2_, pH 7.4]). This and all subsequent steps were performed at 23°C. Cells were fixed for 20 min, washed three times for 5 min in 1 ml sodium cacodylate buffer, and stained with 50 μl 2% OsO_4_ for 1 h. The pellet was washed three times in 1 ml water for 5 min, followed by incubation in 1-2 drops of 4% uranyl acetate solution for 30 min. Stained cells were dehydrated in a graded ethanol series followed by acetone, and embedded in epoxy resin EMBed-812 (Electron Microscopy Sciences). Thin sections (80–100 nm) were cut, post-stained with saturated uranyl acetate for 20 min, rinsed with water, dried, stained with Reynolds’ lead citrate for 10 min, and dried again. TEM images were collected on a JEOL-JEM 1400 Plus transmission electron microscope equipped with a LaB6 filament and operated at an accelerating voltage of 120 kV. Images were recorded using a charge-coupled device camera (Gatan Orius SC1000B).

#### Toxicity assays

For cytotoxicity assays, 1.5 × 10^4^ HEK293T cells were seeded in 96-well plates. After 24 h, cells were transfected with Lipofectamine 2000 (Thermo Fisher Scientific) according to manufacturer’s instructions. Each well received varying amounts of HA-tagged CHMP3 construct, shown to result in equal expression across all constructs by Western blot. After 48 hours, cellular toxicity was analyzed using the CellTiter-Glo Luminescent cell viability assay (Promega) following manufacturer’s instructions. Luminescence was detected using a Biotek Synergy microplate reader.

To confirm equal expression levels, 5 × 10^5^ HEK293T cells were seeded and transfected in parallel in 6-well plates. After 48 hours, cells were harvested, pelleted and lysed in Pierce RIPA Buffer (Pierce) containing Halt Protease Inhibitor (Thermo Fisher Scientific). Samples were mixed with equal amounts 2x Laemmli Sample Buffer containing 10% 2-mercaptoethanol and heated at 95°C for 30 minutes. Samples were run on Mini-Protean TGX™ Precast Protein Gels (Biorad) and transferred to Immobilon-P PDVF Membrane (Millipore). Membranes were blocked in PBS + 0.1% Tween 20 + 5% powdered milk for 1 hour and then incubated with anti-HA (Sigma-Aldrich, H6908) or Anti-Actin monoclonal antibody made in mouse (ThermoFisher Scientific AM4302) for 2 hours at room temperature. After three washes in PBS + 0.1% Tween 20, membranes were incubated in Anti-Rabbit IgG HRP conjugate (Millipore #AP132) or Anti-Mouse IgG/IgM HRP conjugate (Millipore #AP130P) for 2 hours, washed three times in PBS + 0.1% Tween 20 and developed with Advansta WesternBright ECL HRP substrate. Blots were imaged using a LI-COR blot scanner.

#### Immunofluorescence imaging

1 × 10^5^ HeLa N cells were seeded on chambered coverslips (µ-Slide 4 well, ibidi). After 24 h, cells were transfected with Lipofectamine 2000 (Thermo Fisher Scientific) according to manufacturer’s instructions. Each well received varying amounts of HA-tagged CHMP3 construct, once in PBS and fixed in 4% paraformaldehyde for 30 min at room temperature, then washed three times with PBS. Cells were permeabilized with 0.1% Triton-100 for 30 min, then washed three times with PBS. Cells were blocked with sterile-filtered 2% BSA + 0.1% saponin for one hour. Cells were incubated with primary antibody (mouse anti-α-tubulin (Sigma-Aldrich T5168), rabbit anti-HA (Sigma-Aldrich H6908) diluted 1:250 in blocking buffer) overnight at 4°C. The coverslips were washed three times with PBS and incubated in secondary antibody (goat-anti-rabbit Alexa Fluor 488 (Thermo Fisher Scientific A-11034), goat-anti-mouse Alexa Fluor 594 (Thermo Fisher Scientific A-11032) diluted 1:1000 in blocking buffer) for 2 h at room temperature. Cells were washed three times with PBS and covered with DAPI Fluoromount-G (Southern Biotech). Images were captured with a Nikon A1R laser scanning confocal microscope

#### Midbody scoring

To assess abscission defects in unsynchronized cultures, 1 × 10^5^ HeLa N cells were seeded on chambered coverslips (µ-Slide 4 well, ibidi) and 24-well plates 18-24 h before transfection. Cells were transfected with 500 ng empty vector, 25 ng pEF-1alpha-squirrel-monkey-full-length CHMP3-HA, 250 ng pEF-1alpha-squirrel-monkey-CHMP3-HA(155) or 500 ng pEF-1alpha-squirrel-monkey-retroCHMP3-HA using 5 μl Lipofectamine 2000 (Thermo Fisher Scientific) according to manufacturer’s instructions. Total plasmid quantities were equalized using empty vector. Transfection mixes were prepared as 2.5fold master mix and distributed between coverslips and 24-wells. The medium was replaced with 1 ml DMEM 4-6 h later. 20 h post-transfection, cells in 24-well plates were washed once with PBS, detached using 1x TrypLE Express (Gibco, ThermoFisher Scientific) and lysed for 5 min on ice in 50 μl Triton lysis buffer (50 mM Tris pH Aldrich). 50 μl 2× Laemmli SDS-PAGE loading buffer supplemented with 10% 2-mercaptoethanol (Sigma-Aldrich) were added and samples were boiled for 10 min. Proteins were separated by SDS-PAGE, transferred onto PVDF membranes and probed with antibodies to confirm equal expression levels. Cells on coverslips were fixed in 4% paraformaldehyde for 20 min at room temperature and processed for immunofluorescence as described above.

For synchronization experiments, 1 × 10^5^ HeLa N cells were seeded on chambered coverslips (µ-Slide 4 well, ibidi) 18-24 h before transfection. Cells were transfected with 500 ng empty vector, 250 ng pEF-1alpha-squirrel-monkey-CHMP3-HA(155) or 500 ng pEF-1alpha-squirrel-monkey-retroCHMP3-HA using 5 μl Lipofectamine 2000 (Thermo Fisher Scientific) according to manufacturer’s instructions. Total plasmid quantities were equalized using empty vector. 8h after transfection, the medium was replaced with DMEM supplemented with 2mM thymidine (Sigma Aldrich) to synchronize cells. 24 h later, cells were released from thymidine arrest by washing three times with 1x PBS and addition of fresh DMEM. At indicated timepoints, cells on coverslips were fixed in 4% paraformaldehyde for 20 min at room temperature and processed for immunofluorescence as described above.

Images were captured with an Axio Imager M2 (Carl Zeiss) equipped with a Colibri7 LED light source (Carl Zeiss), a pco.edge 4.2 CMOS camera microscope, and a 63X oil-immersion objective (Carl Zeiss, Plan Apochromat, NA 1.4). Laser settings for each channel were confirmed with the untransfected well to ensure no false positives in the green channel. 200 nm optical Z-sections were captured for each field and the z-stack was cycled through. Numbers of midbodies in transfected, HA-positive cells were assessed by a blinded reviewer.

#### EGF receptor degradation

0.6 × 10^6^ HEK293T cells/well were seeded in 6-well plates 24 h before transfection. Cells were transfected with 1 μg empty vector, 1 μg pEF-1alpha-squirrel-monkey-retroCHMP3-HA or 500 ng pGFP-VPS4A E228Q using 10 μl Lipofectamine 2000 (Thermo Fisher Scientific) according to manufacturer’s instructions. Total plasmid quantities were equalized using empty vector. 5-6 h after transfection, media was aspirated and replaced with DMEM without FBS to starve cells. 20h after transfection, starvation media was replaced with DMEM supplemented with 10% FBS, 100 ng/ml recombinant human EGF protein (Thermo Fisher Scientific) and 10 μg/ml cycloheximide (Sigma Aldrich). Cells were harvested at indicated timepoints by detaching in TrypLE Express (Gibco, ThermoFisher Scientific) and lysed in RIPA buffer (50 mM Tris-HCl pH 8.0, 150 mM NaCl, 1% NP-40, 0.5% Na-deoxycholate, 0.1% SDS, 0.1 mM EDTA) supplemented with mammalian protease inhibitor (Sigma-Aldrich) for 15 minutes on ice. The protein concentration in lysates was determined using the Pierce BCA Protein Assay Kit (Thermo Scientific) and concentration was adjusted using RIPA buffer. Equal amounts of 2× Laemmli SDS-PAGE loading buffer supplemented with 10% 2-mercaptoethanol (Sigma-Aldrich) were added, and samples were boiled for 10 min. Proteins were separated by SDS-PAGE, transferred onto nitrocellulose membranes and probed with antibodies. Primary antibodies were as follows: anti-EGF receptor (Cell Signaling Technology, D38B1), anti-tubulin (Sigma Aldrich, T5168), anti-HA (Sigma-Aldrich, H6908), anti-GFP (Roche, 11814460001). Bands were visualized by probing the membrane with goat-anti-rabbit or anti-mouse HRP-conjugated (Sigma Aldrich,) and enhanced chemiluminescence substrate (Thermo Fisher), and detected with a ChemiDoc MP imaging system (Bio-rad). Band intensities were quantified using LI-COR Image Studio software and normalized to band intensities at t = 0h.

#### Pulldown experiments

For pulldown experiments utilizing the One-STrEP-FLAG (OSF) tag, HEK293T cells were seeded in a 6-well plate at 0.8 × 10^6^ cells/well and co-transfected 18–24 h later using 10 μl Lipofectamine 2000 (Thermo Fisher Scientific) with 1.5 µg CHMP4B and the following amounts of CHMP3-expressing plasmids: 1 µg each of pCAG-OSF-PP-human-CHMP3, pCAG-OSF-PP-human-CHMP3(150), pCAG-OSF-PP-squirrel-monkey-CHMP and pCAG-OSF-PP-squirrel-monkey-CHMP3(155) and 3 µg pCAG-OSF-PP-squirrel-monkey-retroCHMP3. Total plasmid quantities were equalized using pCAG-OSF-PP empty vector. The media was changed to fresh DMEM 6-8 h post-transfection. Cells were harvested 24 h post-transfection and lysed in 200 µl RIPA buffer (50 mM Tris-HCl pH 8.0, 150 mM NaCl, 1% NP-40, 0.5% Na-deoxycholate, 0.1% SDS, 0.1 mM EDTA) or NP-40 buffer (10 mM Tris-HCl pH 8.0, 150 mM NaCl, 1% NP-40) supplemented with mammalian protease inhibitor (Sigma-Aldrich) on ice for 15 min. Lysates were clarified by centrifugation at 16,100×*g* for 10 min at 4°C. Cell lysates were diluted to 1 ml with RIPA buffer without SDS or NP-40 buffer and incubated for 45 min at 4°C with 20 µl pre-equilibrated Strep-Tactin Sepharose resin (IBA Lifesciences). The resin was washed three times with RIPA buffer without SDS or with NP-40 buffer. After the final wash, the streptactin beads were aspirated to near dryness, and bound proteins were eluted by boiling in 40 µl 2× Laemmli SDS-PAGE loading buffer supplemented with 10% 2-mercaptoethanol (Sigma-Aldrich), resolved by SDS-PAGE, transferred onto PVDF membranes and probed with primary antibodies (anti-myc (EMD Millipore, 05-724), anti-FLAG (Sigma-Aldrich, F1804). Bands were visualized by probing the membrane with fluorescently labeled secondary antibodies (Li-Cor Biosciences) and scanning with an Odyssey Imager (Li-Cor Biosciences).

#### Live-cell imaging

HeLa and HEK293T cells for live cell imaging were seeded onto MatTek dishes with no. 1.5 coverslips coated with fibronectin (Invitrogen). HeLa cells were transfected 6 hours before imaging with Fugene 6 (Promega) and 1000 ng of DNA. HEK293T cells were transfected with Lipofectamine 2000 (Thermo Fisher) and 1000 ng of DNA 5 h before imaging. Cells were transfected with 600 ng untagged pCRV1-HIV-1NL4.3 GagPol and 150 ng pCRV1-HIV-1NL4.3 GagPol-mEGFP and 250 ng pLNCX-mCherry-CHMP4B or VPS4A. Samples with retroCHMP3 additionally received 250 ng pEF1α-squirrel-monkey-retroCHMP3-T2A-H2B-tagBFP, which encodes contains a self-cleaving H2B-BFP fusion protein for identification of transfected cells.

Live cell imaging was performed with a custom-built microscope that is based on an Olympus IX-81 frame and equipped with a custom-built through-the-objective polarized TIRFM illuminator (Johnson et al., 2014). A 100X PLANAPO 1.50NA Olympus Objective was used. A 488 nm (100 mW LuxX diode laser, Omicron), a 594 nm laser (100 mW iode-pumped solid-state laser, Cobolt AB), and a 405 nm laser (100 mW LuxX diode laser, Omicron) were used at 25 mW, 30 mW, and 2 mW respectively. For live cells, the temperature was maintained at 37°C throughout imaging using custom-built housing for the microscope. TIR light was azimuthally scanned at 100 Hz with mirror galvanometers (Nutfield Technology). Emission was collected sequentially using a Chroma ZET405/488/594m filter and a CMOS camera (Flash-4.0, Hamamatsu). Image acquisition was done with MetaMorph software. All exposure times are 200 ms. GagPol-mEGFP images were taken every 20 s and ESCRT images were taken every 5 s.

HeLa and HEK293T cell assembly videos were analyzed with MetaMorph software. The camera has 100 units added to each pixel and this we subtracted from all frames prior to analysis. The images of GagPol assembly with excitation at 488 nm were analyzed first and all assemblies were marked with circular regions that encompassed the entire assembling particle. Assemblies were confirmed by the shape of the intensity curve over time which shows accumulation and plateau as shown previously (Jouvenet et al., 2008; Jouvenet et al., 2006). After all assemblies were selected, the regions from the assemblies were overlaid onto the ESCRT images taken with excitation at 594 nm of the same cell. Any spikes in intensity from the ESCRT regions were checked to confirm that they correspond to puncta with a Gaussian distribution that overlapped with the GagPol puncta. The number of frames that a single ESCRT puncta stayed and the number of recruitments per assembly was recorded.

To plot assemblies, each assembly was selected via a circular region that encompassed the whole puncta and was trimmed to the first time the assembling puncta was visible and the last time the puncta was visible. This automatically set the first frame of the assembly as time zero. To account for the differing intensities between different cells, the intensities for each assembly trace were rescaled by the following equation: 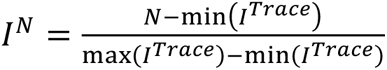. After rescaling, a rolling filter was applied to the traces. The rescaled max intensity for each frame of the assemblies were plotted together using python. For plotting ESCRT recruitment, the same circular region was used as the assembly the ESCRT was recruited to and the trace was trimmed to the same first and last time as the puncta. No filters or rescaling were applied to the ESCRT traces.

To plot overall number of repeat recruitments and length of recruitment, histograms were made which plotted the percent of total counts for either CHMP4B or VPS4A with or without retroCHMP3. Statistical differences between histograms were calculated using a one-tailed T-test. To plot the final gag assembly intensity, the average of the final 3 frames of each trace was plotted. Statistical significance was calculated using a two-tailed T-test.

#### Detection of retroCHMP3 RNA

Total RNA was isolated from cultured squirrel monkey and mouse cell lines according to the manufacturer’s instructions using the Zymo Quick-RNA Miniprep kit (Zymo Research), including the In-column DNAse 1 treatment. For expression analysis in mouse tissues, dissected tissues from male C57BL/6J mice (supplied by the Transgenic Gene-Targeting Mouse Facility at the University of Utah) were homogenized in 1 ml TRIzol reagent (Thermo Fisher Scientific) in a bead homogenizer and RNA was extracted according to manufacturer’s instructions. In order to remove any contaminating genomic DNA, total RNA was treated with TURBO DNase (Thermo Fisher Scientific) prior to cDNA synthesis.

For cDNA synthesis, 2 µg total RNA was reverse transcribed according to the manufacturer’s instructions using the Maxima First Strand Synthesis Kit with DNase treatment (Thermo Fisher Scientific). Thus, each sample underwent two rounds of DNAase treatment for removal of genomic DNA. Additionally, all samples were synthesized in duplicate, minus or plus reverse transcriptase.

PCR amplification of products was performed with Phusion Flash High-Fidelity PCR Master Mix (Thermo Scientific). For each amplification, the -/+ RT samples were treated equally. The minus RT serves as a control to ensure complete removal of genomic DNA from samples; any product in the +RT sample is the result of RNA transcript, as opposed to the –RT control, which indicates genomic DNA contamination. Retrocopies of CHMP3 were amplified 30 to 34 cycles due to their low abundance. Primer sequences used are listed in Table S2. All bands were excised, cloned and sequenced to verify a complete match with the retroCHMP3 sequence.

#### Promoter analysis

The upstream region of the mouse retroCHMP3 sequence was examined for regulatory elements on the USCS genome browser (GRCm38/mm10). The DNA sequence 15kb upstream of the start codon was extracted and run through the NSite (Shahmuradov and Solovyev, 2015) and CiiiDER (Gearing et al., 2019) prediction tools.

#### Interferon stimulation

For interferon stimulation, 2 x 10^6^ mouse cardiac endothelial cells (MCEC) or 6 x 10^6^ squirrel monkey B cells were induced for 6, 12, or 20 h with the addition of 1000 U/mL IFN mix (*e. coli*-derived mouse IFN-gamma protein (R&D Systems, 485-MI-100), HEK293-derived mouse IFN-beta protein (R&D Systems, 8234-MB-010 and Universal Type I Interferon (pbl assay science, 11200-2)), or none as a control. Cells were trypsinized and pelleted, resuspended in RNA lysis buffer; the control flask was collected together with the 6hr IFN treated flask. Pellets were stored at −80°C until RNA extraction. Total RNA was extracted using the Quick-RNA Miniprep Kit (Zymo) and processed as described above. RetroCHMP3, ISG15, and β-actin or CTCF were amplified from the cDNA products (+/-RT) using Phusion Flash High-Fidelity Master Mix (Thermo Scientific). Primers are listed in Table S2.

#### Droplet digital PCR

Freshly synthesized cDNA was frozen at −80°C until submission. Succinyl dehydrogenase (SDHA) was used as a low copy housekeeping transcript and RNaseL was used as a positive control for interferon stimulation (Malathi et al., 2007). In all cases, 11 µL of BioRad ddPCR Supermix for Probes (no dUTP) were mixed with 1.1 µL of specific 20x assays and 9.9 µL cDNA template. For detection of retroCHMP3, 9.9 µL cDNA was added neat. To compare SDHA to retroCHMP3, 9.9 µL of 1:100 dilution was added to the SDHA primer/probe assay. For RNaseL and SDHA duplexes, 9.9 µL of 1:10 dilution was added to the assay. Droplets were generated and PCR was performed using the BioRad QX200 AutoDG Droplet Digital PCR System. A single cycle of 10 min at 95°C was followed by 50 cycles of 95°C for 15 seconds and 60°C for 60 seconds, followed by a single cycle of 98°C for 10 min. Primers and probes are listed in Table S2.

Data were analyzed using BioRad Quantasoft Analysis Pro software. Copies per 20 µL were calculated after manual gating. The fold change in retroCHMP3 or RNaseL expression was calculated for each time point (6, 12 and 20 h) by dividing copies/20 µL at each time point by the number of copies at time 0. Finally, the relative fold change was calculated by dividing the fold change (as described above) by the ratio of SDHA expression at time 6, 12 or 20 by SDHA expression at time 0.

### QUANTIFICATION AND STATISTICAL ANALYSIS

Statistical significance was determined using the GraphPad Prism 8 software package. Statistical tests used, exact values of n, number of replicates and definition of center, dispersion and precision measures for each experiment are noted in the figure legends. All samples were analyzed without blinding, except for phenotype scoring in Fig. 3B and midbody scoring in Fig. 3C and 3D, which were analyzed by a blinded reviewer.

## References

Altschul, S.F., Gish, W., Miller, W., Myers, E.W., and Lipman, D.J. (1990). Basic local alignment search tool. J Mol Biol 215, 403–410.

Babst, M., Katzmann, D.J., Estepa-Sabal, E.J., Meerloo, T., and Emr, S.D. (2002). Escrt-III: an endosome-associated heterooligomeric protein complex required for mvb sorting. Dev Cell 3, 271–282.

Bache, K.G., Stuffers, S., Malerod, L., Slagsvold, T., Raiborg, C., Lechardeur, D., Walchli, S., Lukacs, G.L., Brech, A., and Stenmark, H. (2006). The ESCRT-III subunit hVps24 is required for degradation but not silencing of the epidermal growth factor receptor. Mol Biol Cell 17, 2513–2523.

Bajorek, M., Schubert, H.L., McCullough, J., Langelier, C., Eckert, D.M., Stubblefield, W.M., Uter, N.T., Myszka, D.G., Hill, C.P., and Sundquist, W.I. (2009). Structural basis for ESCRT-III protein autoinhibition. Nat Struct Mol Biol 16, 754–762.

Bartusch, C., and Prange, R. (2016). ESCRT Requirements for Murine Leukemia Virus Release. Viruses 8, 103.

Baumgartel, V., Ivanchenko, S., Dupont, A., Sergeev, M., Wiseman, P.W., Krausslich, H.G., Brauchle, C., Muller, B., and Lamb, D.C. (2011). Live-cell visualization of dynamics of HIV budding site interactions with an ESCRT component. Nat Cell Biol 13, 469–474.

Bendjennat, M., and Saffarian, S. (2016). The Race against Protease Activation Defines the Role of ESCRTs in HIV Budding. PLoS Pathog 12, e1005657.

Bertin, A., de Franceschi, N., de la Mora, E., Maity, S., Alqabandi, M., Miguet, N., di Cicco, A., Roos, W.H., Mangenot, S., Weissenhorn, W., et al. (2020). Human ESCRT-III polymers assemble on positively curved membranes and induce helical membrane tube formation. Nat Commun 11, 2663.

Bianchi, F., and van den Bogaart, G. (2020). Vacuolar escape of foodborne bacterial pathogens. J Cell Sci 134.

Bleck, M., Itano, M.S., Johnson, D.S., Thomas, V.K., North, A.J., Bieniasz, P.D., and Simon, S.M. (2014). Temporal and spatial organization of ESCRT protein recruitment during HIV-1 budding. Proceedings of the National Academy of Sciences 111, 12211–12216.

Brennan, G., Kozyrev, Y., and Hu, S.L. (2008). TRIMCyp expression in Old World primates Macaca nemestrina and Macaca fascicularis. Proc Natl Acad Sci U S A 105, 3569–3574.

Casola, C., and Betran, E. (2017). The Genomic Impact of Gene Retrocopies: What Have We Learned from Comparative Genomics, Population Genomics, and Transcriptomic Analyses? Genome Biol Evol 9, 1351–1373.

Christ, L., Raiborg, C., Wenzel, E.M., Campsteijn, C., and Stenmark, H. (2017). Cellular Functions and Molecular Mechanisms of the ESCRT Membrane-Scission Machinery. Trends Biochem Sci 42, 42–56.

Chuong, E.B., Elde, N.C., and Feschotte, C. (2017). Regulatory activities of transposable elements: from conflicts to benefits. Nat Rev Genet 18, 71–86.

Ciomborowska, J., Rosikiewicz, W., Szklarczyk, D., Makalowski, W., and Makalowska, I. (2013). “Orphan” retrogenes in the human genome. Mol Biol Evol 30, 384–396.

Daugherty, M.D., and Malik, H.S. (2012). Rules of Engagement: Molecular Insights from Host-Virus Arms Races. Annual Review of Genetics 46, 677–700.

Dukes, J.D., Richardson, J.D., Simmons, R., and Whitley, P. (2008). A dominant-negative ESCRT-III protein perturbs cytokinesis and trafficking to lysosomes. Biochem J 411, 233–239.

Eden, E.R., White, I.J., and Futter, C.E. (2009). Down-regulation of epidermal growth factor receptor signalling within multivesicular bodies. Biochem Soc Trans 37, 173–177.

Effantin, G., Dordor, A., Sandrin, V., Martinelli, N., Sundquist, W.I., Schoehn, G., and Weissenhorn, W. (2013). ESCRT-III CHMP2A and CHMP3 form variable helical polymers in vitro and act synergistically during HIV-1 budding. Cellular Microbiology 15, 213–226.

Elde, N.C., and Malik, H.S. (2009). The evolutionary conundrum of pathogen mimicry. Nat Rev Microbiol 7, 787–797.

Elia, N., Sougrat, R., Spurlin, T.A., Hurley, J.H., and Lippincott-Schwartz, J. (2011). Dynamics of endosomal sorting complex required for transport (ESCRT) machinery during cytokinesis and its role in abscission. Proc Natl Acad Sci U S A 108, 4846–4851.

Friedrich, N., Hagedorn, M., Soldati-Favre, D., and Soldati, T. (2012). Prison break: pathogens’ strategies to egress from host cells. Microbiol Mol Biol Rev 76, 707–720.

Garrus, J.E., von Schwedler, U.K., Pornillos, O.W., Morham, S.G., Zavitz, K.H., Wang, H.E., Wettstein, D.A., Stray, K.M., Cote, M., Rich, R.L., et al. (2001). Tsg101 and the vacuolar protein sorting pathway are essential for HIV-1 budding. Cell 107, 55–65.

Gearing, L.J., Cumming, H.E., Chapman, R., Finkel, A.M., Woodhouse, I.B., Luu, K., Gould, J.A., Forster, S.C., and Hertzog, P.J. (2019). CiiiDER: A tool for predicting and analysing transcription factor binding sites. PLoS One 14, e0215495.

George, S.H., Gertsenstein, M., Vintersten, K., Korets-Smith, E., Murphy, J., Stevens, M.E., Haigh, J.J., and Nagy, A. (2007). Developmental and adult phenotyping directly from mutant embryonic stem cells. Proc Natl Acad Sci U S A 104, 4455–4460.

Gottlinger, H.G., Dorfman, T., Sodroski, J.G., and Haseltine, W.A. (1991). Effect of mutations affecting the p6 gag protein on human immunodeficiency virus particle release. Proc Natl Acad Sci U S A 88, 3195–3199.

Guizetti, J., Schermelleh, L., Mantler, J., Maar, S., Poser, I., Leonhardt, H., Muller-Reichert, T., and Gerlich, D.W. (2011). Cortical constriction during abscission involves helices of ESCRT-III-dependent filaments. Science 331, 1616–1620.

Henne, W.M., Stenmark, H., and Emr, S.D. (2013). Molecular mechanisms of the membrane sculpting ESCRT pathway. Cold Spring Harb Perspect Biol 5.

Hesse, M., Raulf, A., Pilz, G.A., Haberlandt, C., Klein, A.M., Jabs, R., Zaehres, H., Fugemann, C.J., Zimmermann, K., Trebicka, J., et al. (2012). Direct visualization of cell division using high-resolution imaging of M-phase of the cell cycle. Nat Commun 3, 1076.

Johnson, D.S., Bleck, M., and Simon, S.M. (2018). Timing of ESCRT-III protein recruitment and membrane scission during HIV-1 assembly. Elife 7.

Johnson, D.S., Toledo-Crow, R., Mattheyses, A.L., and Simon, S.M. (2014). Polarization-controlled TIRFM with focal drift and spatial field intensity correction. Biophys J 106, 1008–1019.

Jouvenet, N., Bieniasz, P.D., and Simon, S.M. (2008). Imaging the biogenesis of individual HIV-1 virions in live cells. Nature 454, 236–240.

Jouvenet, N., Neil, S.J., Bess, C., Johnson, M.C., Virgen, C.A., Simon, S.M., and Bieniasz, P.D. (2006). Plasma membrane is the site of productive HIV-1 particle assembly. PLoS Biol 4, e435.

Jouvenet, N., Zhadina, M., Bieniasz, P.D., and Simon, S.M. (2011). Dynamics of ESCRT protein recruitment during retroviral assembly. Nature Cell Biology 13, 394–401.

Kaessmann, H., Vinckenbosch, N., and Long, M. (2009). RNA-based gene duplication: mechanistic and evolutionary insights. Nature Reviews Genetics 10, 19–31.

Kent, W.J. (2002). BLAT--the BLAST-like alignment tool. Genome Res 12, 656–664.

Kubiak, M.R., and Makalowska, I. (2017). Protein-Coding Genes’ Retrocopies and Their Functions. Viruses 9.

Lata, S., Roessle, M., Solomons, J., Jamin, M., Gőttlinger, H.G., Svergun, D.I., and Weissenhorn, W. (2008). Structural Basis for Autoinhibition of ESCRT-III CHMP3. Journal of Molecular Biology 378, 818–827.

Lehmann, M.J., Sherer, N.M., Marks, C.B., Pypaert, M., and Mothes, W. (2005). Actin- and myosin-driven movement of viruses along filopodia precedes their entry into cells. J Cell Biol 170, 317–325.

Levy, D.N., Aldrovandi, G.M., Kutsch, O., and Shaw, G.M. (2004). Dynamics of HIV-1 recombination in its natural target cells. Proc Natl Acad Sci U S A 101, 4204–4209.

Liao, C.H., Kuang, Y.Q., Liu, H.L., Zheng, Y.T., and Su, B. (2007). A novel fusion gene, TRIM5-Cyclophilin A in the pig-tailed macaque determines its susceptibility to HIV-1 infection. AIDS 21 *Suppl 8*, S19–26.

Liu, W., Zhou, Y., Peng, T., Zhou, P., Ding, X., Li, Z., Zhong, H., Xu, Y., Chen, S., Hang, H.C., et al. (2018). N(epsilon)-fatty acylation of multiple membrane-associated proteins by Shigella IcsB effector to modulate host function. Nat Microbiol 3, 996–1009.

Lopez-Jimenez, A.T., Cardenal-Munoz, E., Leuba, F., Gerstenmaier, L., Barisch, C., Hagedorn, M., King, J.S., and Soldati, T. (2018). The ESCRT and autophagy machineries cooperate to repair ESX-1-dependent damage at the Mycobacterium-containing vacuole but have opposite impact on containing the infection. PLoS Pathog 14, e1007501.

Mackay, D.R., and Ullman, K.S. (2015). ATR and a Chk1-Aurora B pathway coordinate postmitotic genome surveillance with cytokinetic abscission. Mol Biol Cell 26, 2217–2226.

Malathi, K., Dong, B., Gale, M., Jr., and Silverman, R.H. (2007). Small self-RNA generated by RNase L amplifies antiviral innate immunity. Nature 448, 816–819.

Martin-Serrano, J., Perez-Caballero, D., and Bieniasz, P.D. (2004). Context-dependent effects of L domains and ubiquitination on viral budding. J Virol 78, 5554–5563.

Martin-Serrano, J., Zang, T., and Bieniasz, P.D. (2001). HIV-1 and Ebola virus encode small peptide motifs that recruit Tsg101 to sites of particle assembly to facilitate egress. Nat Med 7, 1313–1319.

McCullough, J., Clippinger, A.K., Talledge, N., Skowyra, M.L., Saunders, M.G., Naismith, T.V., Colf, L.A., Afonine, P., Arthur, C., Sundquist, W.I.,, et al. (2015). Structure and membrane remodeling activity of ESCRT-III helical polymers. Science (New York, NY) 350, 1548–1551.

McCullough, J., Frost, A., and Sundquist, W.I. (2018). Structures, Functions, and Dynamics of ESCRT-III/Vps4 Membrane Remodeling and Fission Complexes. Annu Rev Cell Dev Biol 34, 85–109.

Mehra, A., Zahra, A., Thompson, V., Sirisaengtaksin, N., Wells, A., Porto, M., Koster, S., Penberthy, K., Kubota, Y., Dricot, A., et al. (2013). Mycobacterium tuberculosis type VII secreted effector EsxH targets host ESCRT to impair trafficking. PLoS Pathog 9, e1003734.

Mierzwa, B.E., Chiaruttini, N., Redondo-Morata, L., von Filseck, J.M., Konig, J., Larios, J., Poser, I., Muller-Reichert, T., Scheuring, S., Roux, A., et al. (2017). Dynamic subunit turnover in ESCRT-III assemblies is regulated by Vps4 to mediate membrane remodelling during cytokinesis. Nat Cell Biol 19, 787–798.

Mittal, E., Skowyra, M.L., Uwase, G., Tinaztepe, E., Mehra, A., Koster, S., Hanson, P.I., and Philips, J.A. (2018). Mycobacterium tuberculosis Type VII Secretion System Effectors Differentially Impact the ESCRT Endomembrane Damage Response. mBio 9.

Morita, E., Colf, L.A., Karren, M.A., Sandrin, V., Rodesch, C.K., and Sundquist, W.I. (2010). Human ESCRT-III and VPS4 proteins are required for centrosome and spindle maintenance. Proceedings of the National Academy of Sciences 107, 12889–12894.

Morita, E., Sandrin, V., McCullough, J., Katsuyama, A., Baci Hamilton, I., and Sundquist, Wesley I. (2011). ESCRT-III Protein Requirements for HIV-1 Budding. Cell Host & Microbe 9, 235–242.

Moser von Filseck, J., Barberi, L., Talledge, N., Johnson, I.E., Frost, A., Lenz, M., and Roux, A. (2020). Anisotropic ESCRT-III architecture governs helical membrane tube formation. Nat Commun 11, 1516.

Muziol, T., Pineda-Molina, E., Ravelli, R.B., Zamborlini, A., Usami, Y., Gottlinger, H., and Weissenhorn, W. (2006). Structural basis for budding by the ESCRT-III factor CHMP3. Dev Cell 10, 821–830.

Nahse, V., Christ, L., Stenmark, H., and Campsteijn, C. (2017). The Abscission Checkpoint: Making It to the Final Cut. Trends Cell Biol 27, 1–11.

Neil, S.J., Eastman, S.W., Jouvenet, N., and Bieniasz, P.D. (2006). HIV-1 Vpu promotes release and prevents endocytosis of nascent retrovirus particles from the plasma membrane. PLoS Pathog 2, e39.

Newman, R.M., Hall, L., Kirmaier, A., Pozzi, L.A., Pery, E., Farzan, M., O’Neil, S.P., and Johnson, W. (2008). Evolution of a TRIM5-CypA splice isoform in old world monkeys. PLoS Pathog 4, e1000003.

Nguyen, H.C., Talledge, N., McCullough, J., Sharma, A., Moss, F.R., 3rd, Iwasa, J.H., Vershinin, M.D., Sundquist, W.I., and Frost, A. (2020). Membrane constriction and thinning by sequential ESCRT-III polymerization. Nat Struct Mol Biol 27, 392–399.

Nisole, S., Lynch, C., Stoye, J.P., and Yap, M.W. (2004). A Trim5-cyclophilin A fusion protein found in owl monkey kidney cells can restrict HIV-1. Proc Natl Acad Sci U S A 101, 13324–13328.

Olsen, J.C. (1998). Gene transfer vectors derived from equine infectious anemia virus. Gene Ther 5, 1481–1487.

Philips, J.A., Porto, M.C., Wang, H., Rubin, E.J., and Perrimon, N. (2008). ESCRT factors restrict mycobacterial growth. Proc Natl Acad Sci U S A 105, 3070–3075.

Portal-Celhay, C., Tufariello, J.M., Srivastava, S., Zahra, A., Klevorn, T., Grace, P.S., Mehra, A., Park, H.S., Ernst, J.D., Jacobs, W.R., Jr., et al. (2016). Mycobacterium tuberculosis EsxH inhibits ESCRT-dependent CD4(+) T-cell activation. Nat Microbiol 2, 16232.

Raiborg, C., and Stenmark, H. (2009). The ESCRT machinery in endosomal sorting of ubiquitylated membrane proteins. Nature 458, 445–452.

Randall, R.E., Young, D.F., Goswami, K.K., and Russell, W.C. (1987). Isolation and characterization of monoclonal antibodies to simian virus 5 and their use in revealing antigenic differences between human, canine and simian isolates. J Gen Virol 68 ( *Pt 11*), 2769–2780.

Rheinemann, L., Downhour, D.M., Davenport, K.A., McKeown, A.N., Necessary, C.R., Sundquist, W.I., and Elde, N.C. (2021). Recurrent Emergence of an Antiviral Defense through Repeated Retrotransposition and Truncation of CHMP3. bioRxiv, 2021.2004.2027.441704.

Sandrin, V., and Sundquist, W.I. (2013). ESCRT requirements for EIAV budding. Retrovirology 10, 104.

Sayah, D.M., Sokolskaja, E., Berthoux, L., and Luban, J. (2004). Cyclophilin A retrotransposition into TRIM5 explains owl monkey resistance to HIV-1. Nature 430, 569–573.

Schmitt, A.P., Leser, G.P., Morita, E., Sundquist, W.I., and Lamb, R.A. (2005). Evidence for a new viral late-domain core sequence, FPIV, necessary for budding of a paramyxovirus. J Virol 79, 2988–2997.

Schmitt, A.P., Leser, G.P., Waning, D.L., and Lamb, R.A. (2002). Requirements for budding of paramyxovirus simian virus 5 virus-like particles. J Virol 76, 3952–3964.

Scourfield, E.J., and Martin-Serrano, J. (2017). Growing functions of the ESCRT machinery in cell biology and viral replication. Biochem Soc Trans 45, 613–634.

Shahmuradov, I.A., and Solovyev, V.V. (2015). Nsite, NsiteH and NsiteM computer tools for studying transcription regulatory elements. Bioinformatics 31, 3544–3545.

Sherer, N.M., Lehmann, M.J., Jimenez-Soto, L.F., Ingmundson, A., Horner, S.M., Cicchetti, G., Allen, P.G., Pypaert, M., Cunningham, J.M., and Mothes, W. (2003). Visualization of retroviral replication in living cells reveals budding into multivesicular bodies. Traffic 4, 785–801.

Shim, S., Kimpler, L.A., and Hanson, P.I. (2007). Structure/function analysis of four core ESCRT-III proteins reveals common regulatory role for extreme C-terminal domain. Traffic 8, 1068–1079.

Skowyra, M.L., Schlesinger, P.H., Naismith, T.V., and Hanson, P.I. (2018). Triggered recruitment of ESCRT machinery promotes endolysosomal repair. Science 360.

Song, B., Javanbakht, H., Perron, M., Park, D.H., Stremlau, M., and Sodroski, J. (2005). Retrovirus restriction by TRIM5alpha variants from Old World and New World primates. J Virol 79, 3930–3937.

Strack, B., Calistri, A., Craig, S., Popova, E., and Gottlinger, H.G. (2003). AIP1/ALIX is a binding partner for HIV-1 p6 and EIAV p9 functioning in virus budding. Cell 114, 689–699.

Stremlau, M., Owens, C.M., Perron, M.J., Kiessling, M., Autissier, P., and Sodroski, J. (2004). The cytoplasmic body component TRIM5alpha restricts HIV-1 infection in Old World monkeys. Nature 427, 848–853.

Stuchell-Brereton, M.D., Skalicky, J.J., Kieffer, C., Karren, M.A., Ghaffarian, S., and Sundquist, W.I. (2007). ESCRT-III recognition by VPS4 ATPases. Nature 449, 740–744.

Virgen, C.A., Kratovac, Z., Bieniasz, P.D., and Hatziioannou, T. (2008). Independent genesis of chimeric TRIM5-cyclophilin proteins in two primate species. Proc Natl Acad Sci U S A 105, 3563–3568.

Votteler, J., and Sundquist, W.I. (2013). Virus budding and the ESCRT pathway. Cell Host Microbe 14, 232–241.

Wenzel, E.M., Schultz, S.W., Schink, K.O., Pedersen, N.M., Nahse, V., Carlson, A., Brech, A., Stenmark, H., and Raiborg, C. (2018). Concerted ESCRT and clathrin recruitment waves define the timing and morphology of intraluminal vesicle formation. Nat Commun 9, 2932.

Wilson, S.J., Webb, B.L., Ylinen, L.M., Verschoor, E., Heeney, J.L., and Towers, G.J. (2008). Independent evolution of an antiviral TRIMCyp in rhesus macaques. Proc Natl Acad Sci U S A 105, 3557–3562.

Yang, L., Emerman, M., Malik, H.S., and McLaughlin, R.N.J. (2020). Retrocopying expands the functional repertoire of APOBEC3 antiviral proteins in primates. Elife 9.

Yap, M.W., Young, G.R., Varnaite, R., Morand, S., and Stoye, J.P. (2020). Duplication and divergence of the retrovirus restriction gene Fv1 in Mus caroli allows protection from multiple retroviruses. PLoS Genet 16, e1008471.

Yee, J.K., Miyanohara, A., LaPorte, P., Bouic, K., Burns, J.C., and Friedmann, T. (1994). A general method for the generation of high-titer, pantropic retroviral vectors: highly efficient infection of primary hepatocytes. Proc Natl Acad Sci U S A 91, 9564–9568.

Zamborlini, A., Usami, Y., Radoshitzky, S.R., Popova, E., Palu, G., and Gottlinger, H. (2006). Release of autoinhibition converts ESCRT-III components into potent inhibitors of HIV-1 budding. Proc Natl Acad Sci U S A 103, 19140–19145.

Zhu, J.Y., Abate, M., Rice, P.W., and Cole, C.N. (1991). The ability of simian virus 40 large T antigen to immortalize primary mouse embryo fibroblasts cosegregates with its ability to bind to p53. J Virol 65, 6872–6880.

